# Learned Audiovisual Predictability Modulates Depth-Resolved Hippocampal–Cortical Functional Coupling

**DOI:** 10.64898/2026.10.17.759806

**Authors:** Santa Sozzi, Mahdi Enan, Ryszard Auksztulewicz, Caspar M. Schwiedrzik, Lucia Melloni, Fabrizio Esposito, Nicola Vanello, Federico De Martino

## Abstract

Perception reflects the dynamic integration of prior expectations, uncertainty, and incoming sensory evidence, but the neural circuitry supporting these computations remains debated. Current theories implicate coordinated interactions between the hippocampus and neocortex, potentially at the level of hippocampal subfields and cortical layers. Evidence in humans has come primarily from the visual domain, while less is known about how these interactions support learned auditory predictions. Using submillimetre 7T fMRI, we investigated depth-resolved functional coupling between hippocampal subfields and temporal and frontal cortical regions during audiovisual associative prediction. Participants learned deterministic associations between visual cues and auditory syllables, allowing us to compare auditory processing in predictable and unpredictable contexts. Predictability-related connectivity changes showed a spatially structured pattern involving DG/CA3-pSTG and CA1-inferior frontal regions, with significant edge-wise effects resolved across cortical depths. These findings provide fine-grained evidence that learned auditory predictability modulates hippocampal–cortical functional interactions across temporo-frontal systems.

## 1 Introduction

The brain continuously exploits statistical regularities in the environment to generate predictions about upcoming sensory input [1, 2, 3]. Although the neural mechanisms underlying these computational principles remain debated (see e.g. [4, 5, 6]), most theoretical accounts stress the relevance of network interactions in modulating neural gain, routing information or amplifying sensory representations. Generalised predictive processing treats percepts as inferences under internal generative models [7, 8], which are continuously updated through the interaction between predictions and sensory input. In this view, predictive processing is not confined to local computations within individual regions, but relies on coordinated interactions across distributed hierarchical networks, in which top–down predictions and bottom–up prediction-error signals are exchanged [9, 10, 11, 12]. When described at the mesoscopic cortical scale, these hierarchical interactions have been hypothesized to be instantiated in laminar-specific cortical microcircuits [6, 10]. Alternative accounts emphasise the relevance of dendritic processing or propose different laminar origins for predictions and prediction errors [13, 14, 15, 16], but they likewise attribute specialised roles to cortical layers, reflecting the anatomical segregation of feedforward and feedback pathways [17, 18, 19, 20]. In canonical predictive coding, feedback signals conveying predictions and contextual priors originate predominantly from deep cortical layers and target deep and superficial layers of lower-level regions, whereas feedforward prediction errors arise mainly from superficial layers and project to middle layers of higher-order areas ([6, 21]). This laminar organisation predicts depth-dependent signatures of top–down and bottom–up influences, which should vary systematically with context, for example between predictable and unpredictable inputs.

Beyond the neocortex, an increasing body of work assigns to the hippocampus a central role in predictive processes, placing it at the apex of the cortical hierarchy [17, 21]. Owing to its involvement in (rapid) learning, relational inference, and the generation of prospective representations, the hippocampus is thought to acquire arbitrary associations and use them to anticipate upcoming sensory events (see e.g., [22, 23, 24, 25, 26, 27, 28, 29, 30, 31, 32, 33, 34]. Within this framework, hippocampal–neocortical interactions can flexibly shift between modes that prioritise memory retrieval versus model updating, thus linking mnemonic processing and predictive inference within a shared computational view [35, 36]. As a consequence, hippocampal contributions to prediction are expected to be state-dependent [25]. During the early stages of learning, hippocampal responses may be dominated by violations of expectations that drive updating of internal models, whereas after associations have been consolidated, hippocampal activity may increasingly reflect predicted outcomes [25, 37]. These computations align with complementary hippocampal operations such as pattern separation [38, 39] and pattern completion [39] and can be linked to distinct subfields (i.e., DG/CA3 and CA1 - [40, 41, 42, 28, 31]). Together, these observations suggest that context (e.g., predictability) should dynamically shape the functional interactions between hippocampal subfields and the neocortex, consistent with previous evidence that learned predictiveness modulates hippocampal–sensory cortical coupling [43]. Determining whether these interactions are consistent with neocortical mesoscopic models, therefore, represents a critical step toward understanding the neural implementation of predictive perception.

The advent of ultra-high-field (UHF) MRI has made laminar and depth-resolved functional imaging (fMRI) increasingly feasible, enabling the investigation of mesoscale functional organisation [44, 45, 46], and layer-dependent neocortical interactions in vivo [47, 48, 49]. By providing increased signal-to-noise ratio (SNR) and contrast-to-noise ratio (CNR), UHF imaging has enabled the characterisation of depth-dependent effects related to predictive processes within the neocortex [50, 51, 52, 53, 54] and hippocampal responses related to prediction errors [55], as well as hippocampal-neocortical interactions supporting predictive computations in the visual domain [24]. Despite these advances, our knowledge of how cortical layers participate in the processing of predictive information remains relatively underexplored in domains beyond visual processing. Some works have proved the involvement of broad neocortical (and subcortical) networks for processing auditory predictions (see e.g., [10, 56, 57]). Recent UHF fMRI studies have further revealed layer-dependent responses associated with processing of predictive acoustic information in individual cortical regions [58, 59, 60]. However, while useful for identifying and characterising the regional patterns of local mesoscopic information processing related to expectation, studies targeting isolated regions provide a limited window into the neural communication that enables perceptual inference [10, 61]. Accordingly, evidence of layer-specifc connectivity underlying predictive auditory processes in humans has remained largely indirect or has been inferred primarily from computational modelling of non-invasive electrophysiological recordings [62, 63]. In particular, it remains unclear whether and how activity functionally couples between hippocampal subfields and layers of neocortical regions and is modulated during the processing of auditory contextual information. Closing this gap is especially relevant given the hypothesised role of the hippocampus in predictive inference [21] and its well-established contribution to auditory associative processing [64].

In this work, we leveraged submillimeter 7T laminar fMRI and a cross-modal associative learning paradigm in which auditory content could either be predicted from learned visual cues or remain unpredictable despite matched sensory stimulation. We focused on examining the synchronisation of the functional activity between hippocampal subfields and depth-resolved temporal and prefrontal cortical regions involved in task-relevant auditory prediction and higher-order inference. We investigated how the functional connectivity between hippocampal subfields and neocortical ROIs was shaped by the experimental condition. Specifically, we hypothesised that, if context is expressed in the interaction among areas of the network, then predictability should reorganise functional coupling between the hippocampus and neocortical targets. To isolate contextual (predictive) effects, we contrasted functional connectivity (estimated using beta series correlation) between predictable and unpredictable trials. Specifically, by combining trial-level functional connectivity with laminar fMRI, we investigated if predictability is associated with depth-dependent changes in hippocampal–neocortical functional connectivity. More broadly, this work aims to clarify where (temporal vs. prefrontal systems), across which cortical depths, and under which contextual conditions (predictable vs. unpredictable) hippocampal–cortical interactions are modulated during predictive processing in an associative cross-modal setting, extending laminar evidence beyond classical visual paradigms and towards a mesoscale network-level characterisation of learned associative auditory expectations.

## Results

In a behavioural training session, we verified that participants (N=11) successfully learned the visual–auditory associations underlying the task (Supplementary Figure 4). The same participants, who subsequently underwent fMRI scanning, viewed sequences of images in which a face deterministically predicted an auditory syllable (*PA* or *GA*), enabling the formation of auditory expectations. Unpredictable trials, for which no consistent face-syllable mapping was present, served as a control condition. We observed a significant block (first vs. last) *×* context (predictable vs. unpredictable) interaction (F(1,10) = 18.06, p = 0.0016), with a larger reduction in reaction times for predictable than for unpredictable trials. Post hoc analyses confirmed significant reductions in reaction time across training in both predictable (t(10) = 6.18, p = 0.0001) and unpredictable trials (t(10) = 4.67, p = 0.0009) with accuracy nearing ceiling (mean 99.9%, SD 0.04%). This selective speeding indicates that participants rapidly acquired the face-syllable contingencies and used this information to anticipate the upcoming syllable in the predictable context.

Submillimeter (0.8 mm isotropic) BOLD fMRI data were acquired using gradient-echo EPI over two sessions. Trials were presented in either a predictable or an unpredictable context. In the predictable context, 75% of trials ended with the expected (valid) syllable, whereas the remaining trials presented an invalid syllable [12.5%], or an omission [12.5%]. The unpredictable context also included 12.5% omissions to ensure comparable stimulus statistics. We estimated single-trial fMRI responses to both the visual context (picture) and the syllable sound using GLMsingle [65] (Figure 1 top). Subsequent analyses focused on the auditory portion of each trial.

**Figure 1:**
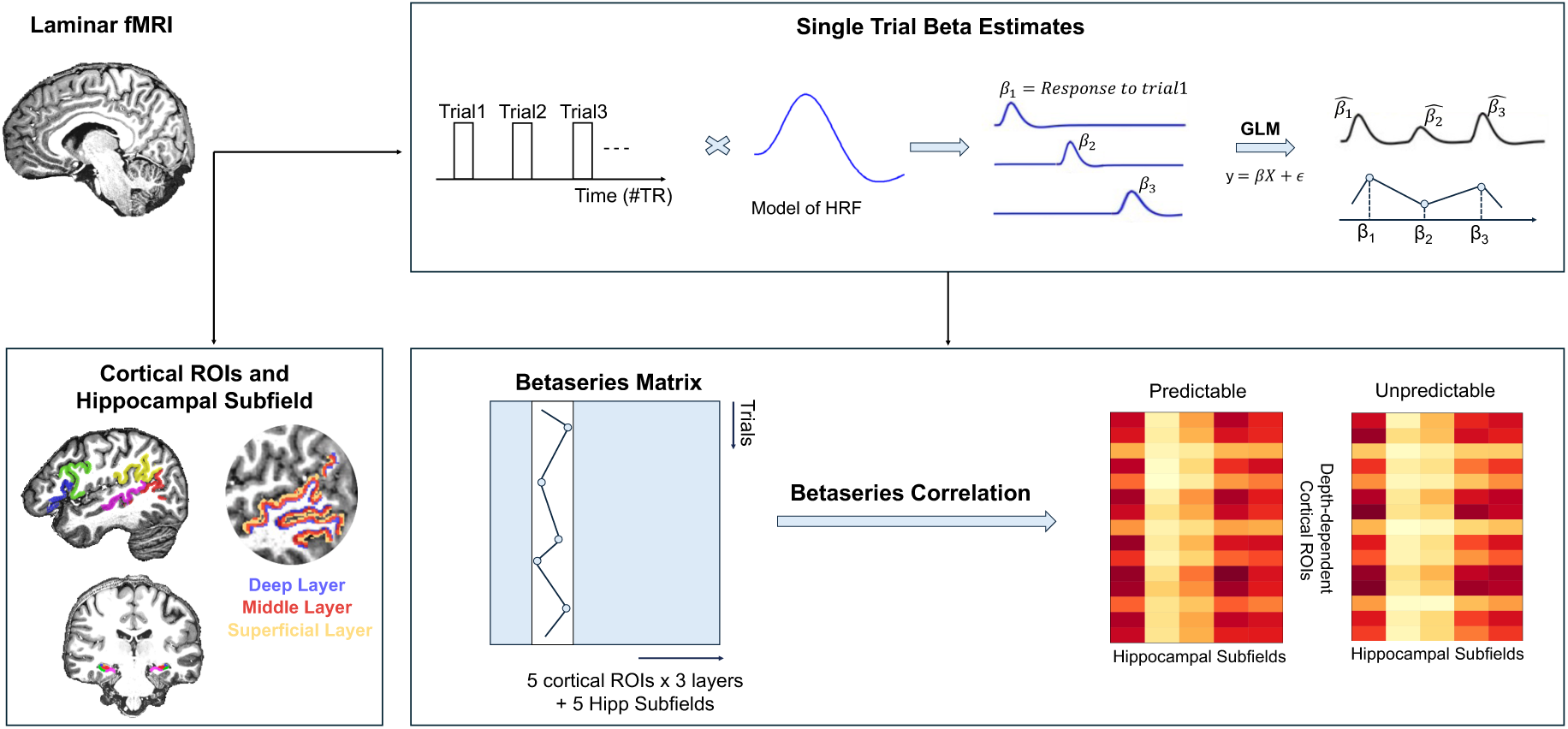
Analysis pipeline. *(Top)* The response to single trials in preprocessed fMRI data was estimated using GLMsingle [65]. *(Bottom left)* Cortical regions of interest were defined on each participant’s cortical surface, projected back into the native volumetric space, and intersected with cortical depths (deep, middle, and superficial) estimated using the equi-volume approach [66, 67]. *(Bottom right)* Beta series were extracted for predictable (valid only) and unpredictable (sound) trials in each cortical layer and hippocampal subfield, resulting in (5 cortical ROIs *×* 3 layers + 5 hippocampal subfields) *×* (number of trials per context, predictable and unpredictable) matrix. Functional connectivity was estimated using beta series correlations between the 5 hippocampal subfields and the 15 layer-specific neocortical ROI. Group-level differences between predictable and unpredictable conditions were assessed using permutations and corrected for multiple comparisons using false discovery rate (FDR).

We defined regions of interest (ROIs) in temporal and frontal cortices using macro-anatomical landmarks on the individual participants’ cortical surfaces. Cortical depths (deep, middle, and superficial) were estimated using the equi-volume approach [66, 67] (Figure 1 bottom left). In a preliminary analysis, we investigated whether the learned task contingencies resulted in cortical depth-dependent validity effects by comparing valid and invalid trials in the predictable context. We sampled percent signal change for each condition, ROI, and cortical depth in each participant, and we averaged across hemispheres, as no hemispheric differences were hypothesized.

The preliminary univariate analyses revealed, as expected, a gradual increase in percent signal change from deep to superficial cortical depths (see Supplementary Figure 5). Moreover, to assess whether the effect of validity varied across layers we performed a non-parametric [Friedman] test for the cortical-depth *×* condition interaction (see Supplementary Table 1 and Supplementary Figure 5). The results identified the posterior superior temporal gyrus [pSTG], posterior middle temporal gyrus [pMTG], pars orbitalis, and pars triangularis as cortical ROIs showing significant depth-dependent validity effects (Figure 1, bottom left panel). All following analyses focused on these cortical ROIs with the addition of the region broadly covering the temporo-parietal junction and the supra marginal gyrus [TPOj], that has been previously implicated in the processing of predictable auditory content in the same paradigm [62]. Individual cornu ammonis (CA1, CA2 and CA3), dentate gyrus (DG) and subiculum (Sub) ROIs were defined in the native anatomical space of each participant using the Hippocampal Segmentation Factory toolbox [68] (Figure 1, bottom left panel; see Supplementary Figure 6 for all individual hippocampal ROIs).

In both neocortical regions and hippocampal subfields, we extracted beta series (i.e., vectors of single-trial BOLD response amplitude estimates to auditory trials, expressed as percent signal change) for predictable (valid only) and unpredictable (sound) contexts. In neocortical ROIs, voxels-wise beta series were averaged within each cortical depth, yielding one representative beta series for the deep, middle, and superficial layers of each ROI. Similarly, voxel-wise beta series were averaged within each hippocampal subfield to obtain a representative beta series for each subfield (Figure 1 bottom left). Hippocampal-cortical functional connectivity was estimated, within each participant, separately for the predictable and unpredictable contexts using beta series correlation (BSC - Figure 1 bottom right). Group-level inference on predictability was based on the within-subject edge-wise difference in Fisher *z*-transformed connectivity (Δ*z*_BSC_ = *z*_predictable_ *− z*_unpredictable_), tested using permutations and corrected for multiple comparisons using false discovery rate. For visualisation, results were presented in matrix form as differences in beta-series correlation coefficients (Δ_BSC_ = BSC_predictable_ *−* BSC_unpredictable_), with positive values indicating stronger coupling in the predictable context and negative values indicating stronger coupling in the unpredictable context.

Differences between the predictable and unpredictable conditions revealed sparse predictability-related effects across the tested hippocampal-neocortical connections (Figure 2A). Significant edge-wise predictability effects involved pSTG-DG/CA3 and inferior frontal-CA1 connectivity, with effects resolved across cortical depths.

In particular, predictability resulted in increased coupling between DG and deep (Δ_BSC_ = 0.1069, *q*_FDR_ = 0.0458) and superficial (Δ_BSC_ = 0.1098, *q*_FDR_ = 0.0488) layers of pSTG, while CA3 showed significant effects with middle (Δ_BSC_ = 0.0685, *q*_FDR_ = 0.0419) and superficial (Δ_BSC_ = 0.0875, *q*_FDR_ = 0.0419) layers of pSTG in the predictable compared with the unpredictable context. We also found significant effects on the modulation of connectivity between the inferior frontal cortical regions (pars orbitalis and pars triangularis) and CA1. Predictability-related connectivity increases with CA1 were observed across all three depths of pars orbitalis (deep: Δ_BSC_ = 0.1104, *q*_FDR_ = 0.0419; middle: Δ_BSC_ = 0.1164, *q*_FDR_ = 0.0419; superficial: Δ_BSC_ = 0.1008, *q*_FDR_ = 0.0419), as well as at middle (Δ_BSC_ = 0.0744, *q*_FDR_ = 0.0419) and superficial (Δ_BSC_ = 0.1045, *q*_FDR_ = 0.0419) layers of pars triangularis. Individual values in the modulation of functional connectivity together with group means and confidence 95% intervals are shown in Figure 2B. All effect sizes and corrected significance values are reported in Supplementary Tables 2-6 for both significant and not significant couplings. In the Supplementary Material, we also report the unthresholded beta-series correlation matrices for the predictable and unpredictable contexts separately, as well as an analysis conducted at the level of whole cortical regions (without depth-dependent separation). The latter highlighted a broadly comparable spatial pattern of predictability-related coupling between hippocampal subfields and neocortical regions when cortical depth was not explicitly resolved.

**Figure 2:**
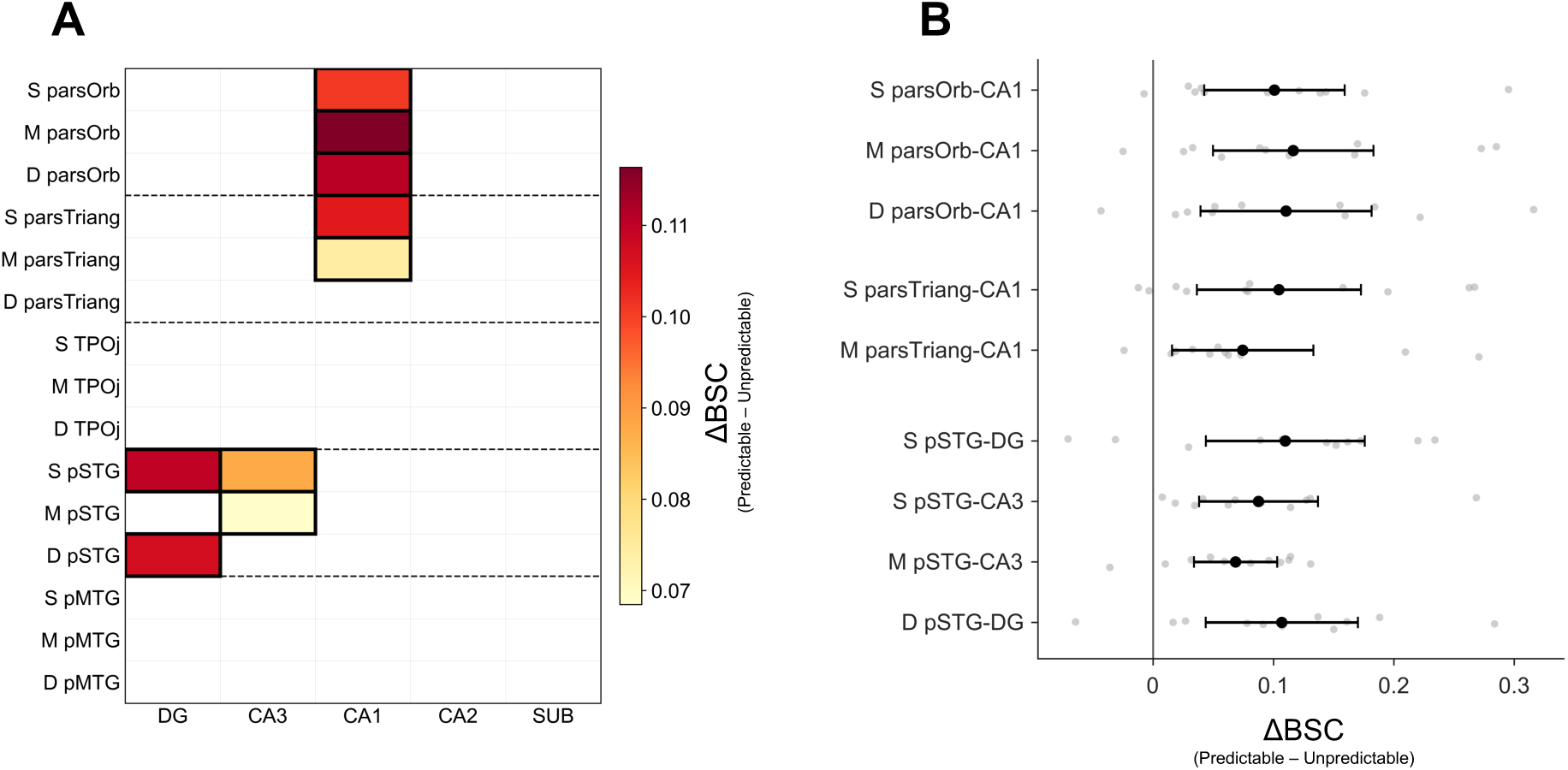
Predictability-related changes in depth-resolved hippocampo–neocortical connectivity. (*A*) Group-level heatmap summarising significant predictability effects on beta-series connectivity for each hippocampal subfield (columns: DG, CA3, CA1, CA2, SUB) and each cortical ROI *×* cortical depth (rows: (S), (M), (D) for Pars Orbitalis, Pars Triangularis, TPOj, pSTG, and pMTG). Values represent the differential connectivity contrast (predictable – unpredictable), shown only for edges surviving non-parametric sign-flipping permutation testing with FDR correction (q(FDR)*<*0.05). (*B*) Individual participant values (grey dots) are shown for the difference in beta-series correlation between predictable and unpredictable conditions. Black dots indicate group means and horizontal error bars correspond to the 95% confidence intervals. Only edges surviving non-parametric sign-flipping permutation testing with FDR correction (*q*_FDR_ *<* 0.05) are shown. S, M, and D indicate superficial, middle, and deep cortical depths, respectively.

## Discussion

The functional coupling between the hippocampus and the neocortex is central to theories explaining the brain capacity to exploit regularities in the environment and to generate predictions about upcoming sensory input. While the literature on the putative roles of hippocampal subfields in the generation of memory-driven predictions and errors is extensive (for a recent review, see e.g., [40]), and neocortical layers are increasingly shown to support predictive perception (see e.g., [13, 14, 15, 16]), less is known about whether (and how) the functional connectivity between hippocampal sub-fields and neocortical layers is modulated to support acoustic perception (but see e.g., [55] for information in the visual domain). Using submillimeter 7T laminar fMRI, we investigated how learned predictability modulates hippocampal–neocortical functional connectivity during audiovisual associations. Predictability-related edge-wise effects involved CA1 coupling with inferior frontal cortical regions and DG/CA3 coupling with posterior superior temporal cortex (Figure 3). Notably, our analyses revealed layer dependent neocortical effects. These findings support the idea that predictive auditory processing relies on structured hippocampal–cortical connectivity across temporofrontal systems [40, 21, 10], whose coupling is modulated by the predictive structure of the task.

**Figure 3:**
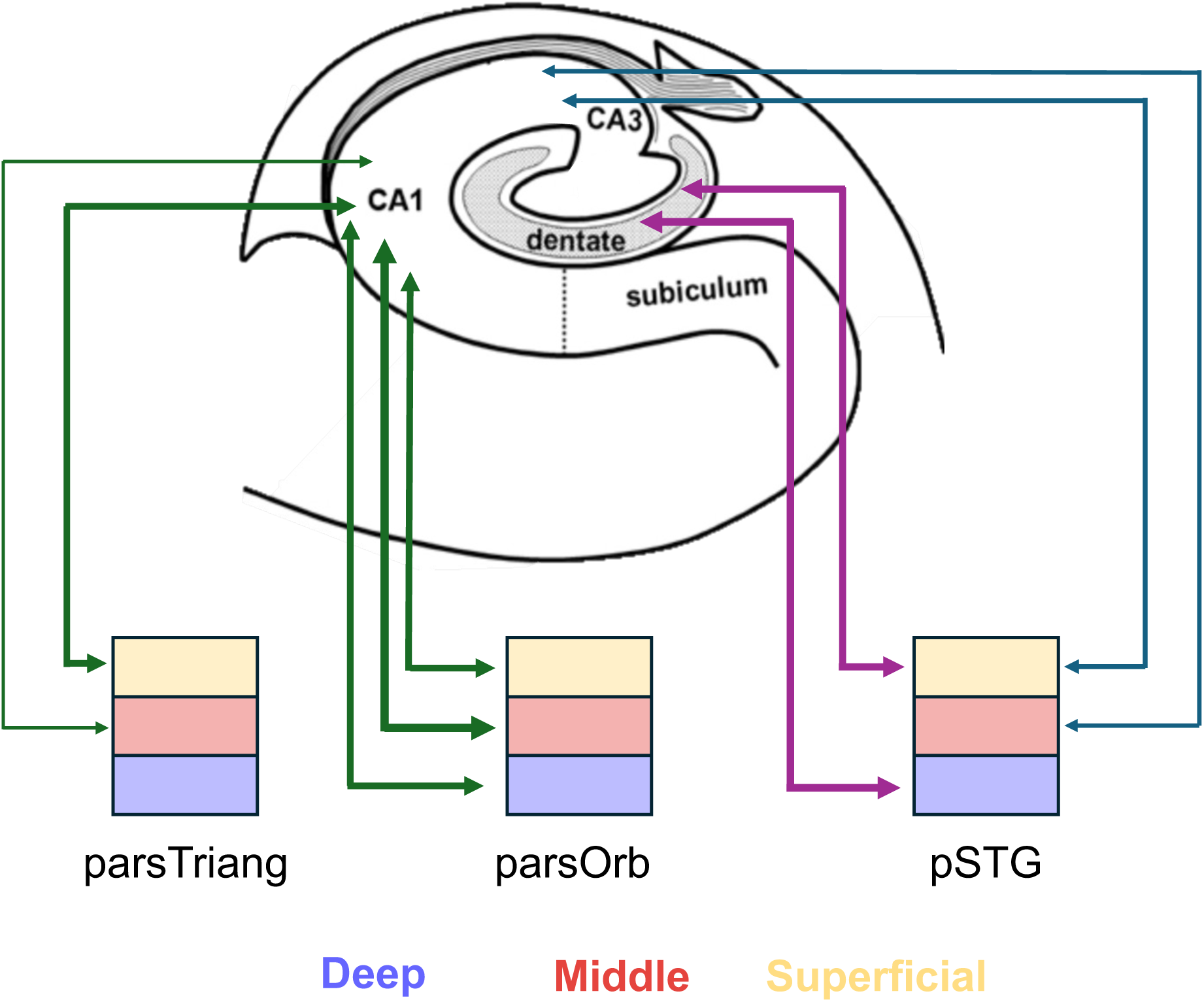
Schematic view of the main hippocampal-neocortical effects. CA1 showed increased coupling with inferior frontal regions (pars Orbitalis and pars Triangularis; green connections), whereas DG (purple connections) and CA3 (blue connections) showed increased coupling with pSTG. Coloured cortical compartments indicate the cortical depths in which significant effects were observed: deep (violet), middle (red), and superficial (yellow). Connections are shown as bidirectional arrows to indicate functional coupling without implying directionality of information flow. Line thickness is proportional to the magnitude of the predictability-related connectivity difference (Δ_BSC_).

Our results revealed predictability-related hippocampal–neocortical connectivity, with significant edge-wise effects involving DG/CA3 and pSTG, and CA1 and inferior frontal cortex. This pattern is consistent with the possibility that CA1 and DG/CA3 contribute differently to predictive processing through their interactions with distributed cortical systems. Increasing evidence suggests that predictive inference emerges through interactions between the hippocampus and distributed cortical networks (see e.g., [34, 22, 43, 24]) and supports the view of the hippocampus as fundamentally relevant to predictive processes [21]. Within this view, hippocampal subfields have been proposed to support partly distinct computations relevant to prediction [30, 69]. In particular, DG and CA3 are thought to contribute to the separation of representations belonging to the same context, with DG supporting particularly fine-grained pattern separation [40]. The recurrent circuitry of CA3 has been implicated in retrieving stored representations from partial cues through pattern completion [39, 70]. On the other hand, CA1 occupies a strategic position in which inputs from CA3 converge with direct entorhinal projections [35]. This convergence has been proposed to allow CA1 both to represent predictive information, particularly as learning progresses and depending on task relevance and relational structure [40], and to compare internally generated predictions with ongoing sensory input [35, 30], thereby contributing to the detection of mismatches between expected and observed information [32].

The increased DG/CA3–pSTG coupling highlighted by our results is consistent with the involvement of hippocampal associative mechanisms in auditory processing according to learned contextual contingencies, and may reflect coordination between hippocampal associative representations and auditory cortical processing during the anticipation of upcoming auditory content. The involvement of posterior STG aligns with its established role in predictive processing of auditory input [63, 71], including invasive electrophysiological evidence gathered using the same design [62]. This interpretation is also consistent with previous work that has shown that hippocampal activity can reinstate sensory representations during memory retrieval and prediction in visual cortex [34, 22], suggesting that similar hippocampal interactions may extend to the auditory domain. More broadly, the present findings are consistent with the possibility that predictive context modulates auditory processing through interactions with hippocampal associative circuitry, in line with models in which recurrent CA3 circuitry contributes to pattern completion and to the stabilisation and updating of associative representations across correlated experiences [72, 70].

The CA1–frontal coupling highlighted by our results may be interpreted in the context of higher-order contextual and relational processing during predictable trials. [40]. Pars orbitalis and pars triangularis have been implicated in contextual integration, controlled retrieval, and selection of task-relevant representations, especially when behaviour depends on using contextual information to guide interpretation or prediction [73, 74]. Increased CA1–frontal coupling in predictable relative to unpredictable contexts may therefore reflect coordination between hippocampal relational representations and frontal mechanisms involved in maintaining and integrating the learned predictive structure of the task. This interpretation is compatible with proposals that state that CA1 contributes to representing relational and predictive information [30, 40], while inferior frontal regions are thought to maintain and integrate task-relevant contextual representations. Although CA1 has also been proposed to act as a comparator between mnemonic expectations and current perceptual input, the present analysis, which focused on valid predictable trials, does not directly test prediction-error or mismatch-related processes. More generally, the observed CA1–frontal coupling may therefore reflect the engagement of higher-order contextual representations during successful prediction, rather than a process specifically related to evaluating prediction validity.

Hippocampal–neocortical coupling was resolved across cortical depths, with significant edge-wise predictability effects observed at particular layers within temporal and frontal regions. The beta series correlation approach does not allow inference regarding the directionality between hippocampal and cortical regions, while cortical-depth BOLD profiles cannot be mapped one-to-one onto underlying synaptic or microcircuit mechanisms because of vascular contributions and signal mixing across cortical layers [75, 76].Contrasting predictable and unpredictable conditions reduces the contribution of depth-dependent vascular biases that are shared across conditions, although residual effects cannot be excluded [77]. Within these constraints, the depth-resolved connectivity patterns provide a spatially finer description of the hippocampal–cortical network engaged during prediction. The occurrence of significant effects at superficial and deep depths in pSTG may be compatible with contextual modulation of auditory representations. Similarly, the broader CA1–frontal interactions may reflect distributed higher-order inferential and contextual computations operating across multiple cortical depths.

Because beta series correlation captures trial-wise covariation in response amplitude, the significant couplings observed in our study reflect statistical dependencies in task-related activity that may be mediated through direct or indirect interactions within the broader network. They should not be interpreted as evidence for direct anatomical projections between hippocampal subfields and neocortical targets. Anatomical tracing studies in non-human primates nevertheless provide a plausible structural framework for these interactions. Direct hippocampal efferents to temporal and prefrontal cortices arise predominantly from CA1, prosubiculum, and subiculum [78, 79], whereas auditory cortical information can reach the hippocampal formation through temporal, perirhinal/parahippocampal, and entorhinal relays that provide input to DG/CA3 as well as CA1 [80, 81, 82]. Parahippocampal regions also project to posterior auditory association cortex [83], providing potential routes through which hippocampal and auditory cortical processing may interact. Accordingly, the DG/CA3–pSTG effects observed here would likely involve indirect pathways if they reflect anatomical communication, whereas CA1–frontal coupling is compatible with both direct and multisynaptic medial-temporal–prefrontal routes [79, 84]. Importantly, however, macaque cortical territories and their projection patterns cannot be mapped one-to-one onto the human cortical ROIs and depth compartments examined here.

Beyond covariation in response amplitude, as we analysed here, establishing whether predictability also modulates the informational content represented within each region would require complementary multivariate approaches, such as representational or informational connectivity analyses [25]. We also acknowledge that the sample size remains modest for connectivity analyses, although it is comparable to that of several previous ultra-high-field laminar fMRI studies, and therefore our results may require replication in independent samples. Finally, beta-series correlation does not establish the temporal order or causal direction of the observed interactions. The depth-resolved hippocampal–cortical connectivity observed here may nevertheless be compatible with recurrent exchanges across hippocampal and cortical systems, a hypothesis that could be tested more directly in future studies using effective-connectivity approaches or generative models of (laminar) BOLD dynamics [85, 59] extended to inter-regional connectivity.

Together, our findings indicate that learned predictive context shapes auditory processing and is associated with changes of hippocampal–neocortical functional coupling within a distributed temporo-frontal network.

## Conclusion

Using submillimetre fMRI to study functional coupling between hippocampal subfields and neocortical depths, we reveal a spatially structured pattern of predictability-related connectivity involving pSTG–DG/CA3 and inferior frontal–CA1 interactions. These results provide a characterisation of the spatial organisation of hippocampal–cortical coupling during learned auditory prediction. More broadly, these results contribute to growing evidence for hippocampal involvement in predictive processing by showing that learned relational knowledge modulates changes in functional coupling between hippocampal subfields and neocortical systems involved in auditory and contextual processing.

## Methods

### 1.1 Participants

Sixteen healthy volunteers (10 females, 6 males; mean age = 25.6 *±* 3.8 years) participated in the study and gave written informed consent prior to MRI scanning. All participants reported no history of neurological or hearing disorders and had normal or corrected-to-normal vision. The study was approved by the Ethics Review Committee for Psychology and Neuroscience (ERCPN) at Maastricht University and was conducted in accordance with the Declaration of Helsinki. Five participants were excluded from the final analyses due to large position differences (identified via visual inspection) across the two sessions (with resulting differences in distortions) that could not be corrected (two participants), the participants’ inability to complete the task (two participants), and head coil size restrictions (one participant). The final sample therefore consisted of eleven participants (8 females, 3 males; mean age = 25.6 *±* 3.7 years).

### 1.2 Experimental Design

Participants performed a visual-auditory associative learning task designed to manipulate the predictability of auditory stimuli based on preceding visual cues. The experimental design was adapted from [62] and consisted of a behavioural training session and an experimental session. Compared to [62] we focused exclusively on content-based (temporally regular) predictions in an event-related design adapted to fMRI.

In the behavioural training, the same participants who subsequently underwent fMRI scanning viewed sequences of images in which a face deterministically predicted an auditory syllable (*PA* or *GA*), enabling the formation of auditory expectations. Participants were presented with trials in which a sequence of visual stimuli (a noise image [1 s] consisting of horizontal and vertical grey-scale patterns, followed by a scene [1 s] and a face [1 s]) was followed by an auditory syllable (*PA* or *GA*, 300 ms). Stimulus categories always occurred in the same temporal order, while the specific scene and face exemplars varied across trials. Participants completed at least four behavioural runs containing 100 sequences each. Trials belonged to one of two types: predictable and unpredictable. In predictable trials, the entire stimulus sequence was deterministic, with the scene predicting the identity of the subsequent face with 50% probability and the face deterministically predicting the following auditory syllable (100% contingency). The unpredictable condition, in which there was no predictive structure (i.e., no consistent face-syllable mapping), served as a control condition. Each face was drawn from eight possible exemplars and each scene from four, with separate stimulus sets for predictable and unpredictable sequences. During training, participants reported, as quickly and accurately as possible, which syllable they heard (*PA* vs. *GA*) via a button press. When responses were slower than 1.5 seconds, participants were prompted to respond faster, and missed trials were excluded from the behavioural analysis. Reaction times and accuracy were recorded for all trials. Predictable and unpredictable trials were randomly intermixed, with a balanced presentation of *PA*/*GA* syllables across behavioural runs.

During MRI acquisition, participants performed the same associative task over two scanning sessions (approximately 84 minutes each). The predictable and unpredictable conditions were presented in separate functional runs. In predictable runs (eight in total) we presented 24 trials per run, and each trial ended with either the expected (valid) syllable (75%), an invalid syllable (12.5%) or an omission (12.5%) (Supplementary Figure 9).

In unpredictable runs (six in total) we presented 16 trials per run, with 12.5% of these trials being omissions to ensure comparable stimulus statistics (Supplementary Figure 9). The timing of visual pictures and auditory syllables was kept identical to that used during behavioural training, but the participant responses were delayed and cued to minimise motor contamination of the auditory responses (Supplementary Figure 9). Participants were instructed to report which syllable they heard or, in omission trials, which syllable they believed should have been presented using a button for the syllable “PA” and a separate button for the syllable “GA”. The experimental design followed a slow event-related design. The response cue appeared 7.5 or 10 seconds after syllable onset (3 or 4 TRs) and was followed by an additional delay of 3–4 TRs before the next trial (randomised across trials; see Supplementary Figure 9). Syllables were presented via MRI-compatible earbuds (Sensimetrics, model S15) at comfortable listening levels that were individually adjusted to ensure audibility over scanner noise. When no visual stimulus or cue was shown, a fixation cross remained on the screen. Across the two sessions, each participant completed 288 trials in total (192 predictable and 96 unpredictable).

### 1.3 MRI data acquisition

We collected data using a 7 T whole-body MRI system (Siemens Healthineers, Erlangen, Germany) equipped with a 32-channels receive head coil (single transmit; Nova Medical Inc., Wilmington, MA, USA). The acquisition protocol included a high-resolution T1-weighted anatomical image (MP2RAGE, 0.7 mm isotropic resolution, 240 slices, TR = 5000 ms, TE = 2.39 ms, TI_1_/TI_2_ = 800/2750 ms, FA_1_/FA_2_ = 4*^◦^/*5*^◦^*, GRAPPA acceleration factor = 3) and functional T2*^∗^*-weighted images acquired using a multiband gradient-echo echo-planar imaging (GE-EPI) sequence (0.8 mm isotropic resolution, 66 slices, TR = 2500 ms, TE = 26 ms, multiband factor = 2, GRAPPA acceleration factor = 3). Functional images covered frontal, temporal, and occipital regions bilaterally. For each participant, five reverse phase-encoded volumes were also acquired to estimate susceptibility-induced distortions and perform distortion correction during preprocessing. All data are available at https://openneuro.org/datasets/ds007109.

### 1.4 Behavioural Analysis

Behavioural performance during training was assessed using reaction times and response accuracy. To evaluate learning across training and its dependence on context, reaction times were analysed using a block (first vs. last) *×* context (predictable vs. unpredictable) analysis. Significant interaction effects were further examined using post hoc comparisons within each context. Response accuracy was summarised descriptively across participants.

### 1.5 fMRI Data Analysis

BrainVoyager (v22.2, Brain Innovation, Maastricht, The Netherlands) [86] was used in all anatomical and functional preprocessing steps unless otherwise specified. The preprocessing pipeline developed for this study, together with the custom analysis code, is available online (https://github.com/mesoScopic-Computational-AuditioN-lab/ResearchProjects/tree/main/finalized/AssociativeLearning)

#### 1.5.1 (f)MRI preprocessing

The unbiased T1-weighted image (MP2RAGE) was denoised [87], skull-stripped using Nighres [88], corrected for residual intensity inhomogeneities, and upsampled to an isotropic resolution of 0.4 mm. Up-sampling was necessary because, in BrainVoyager, functional data are analysed at a resolution that is an integer multiple of the resolution of the anatomical data. By up-sampling the anatomical data to 0.4 mm isotropic resolution, we could maintain the native functional resolution of 0.8 mm isotropic. The cortical grey matter ribbon was segmented using the built-in 3D patchwise Tiramisu deep-learning network for segmentation of submillimetre resolution 7T brain images implemented in BrainVoyager [89]. The resulting segmentation was visually inspected and manually corrected where necessary using ITK-SNAP [90]. Corrections were necessary in (portions of the) temporal lobes where the most typical automatic segmentation error was an underestimation of the thickness of the grey matter. Cortical layers were subsequently generated within the gray matter ribbon using an equi-volume model, yielding deep, middle, and superficial layers.

Functional images were preprocessed using slice-scan-time correction, motion correction (using the first volume of the first run of the first session as reference) and high-pass filtering. The correction of susceptibility-induced distortions was performed using the reversed phase-encoding data with Topup (FSL v6.0.5). When necessary, residual distortions were further corrected using Advanced Normalization Tools (ANTs; antsRegistration), including both linear and non-linear transformations. When the standard motion correction procedure failed (i.e., when visual inspection revealed incorrect alignment for [part of] the run), we first aligned all functional volumes of a given run to the first volume of the same run, then averaged the first five volumes and aligned these averages across runs (the first run of session one as reference) to obtain between-runs motion parameters. Note that the combined motion parameters of both within-run and between-run alignment were re-applied to the original time series to minimise the number of interpolation steps. Finally, functional images were registered to the anatomical images using boundary-based registration (BBR) [91].

#### 1.5.2 ROI definition

We defined nine regions of interest (ROIs) on the individual mid gray matter surface, separately for the two hemispheres. In particular, following macro-anatomical landmarks (gyri and sulci) [92, 93], we divided the temporal lobe into Heschl’s Gyrus (HG), Planum Polare (PP), Planum Temporale (PT), anterior Superior Temporal Gyrus (aSTG), posterior Superior Temporal Gyrus (pSTG). In addition, we defined two regions in the frontal cortex bilaterally, the pars Orbitalis and pars Triangularis, one region covering the posterior Middle Temporal Gyrus (pMTG) and one region broadly covering the Temporal-Parietal-Occipital junction and the Supra Marginal Gyrus (TPOj). ROIs were projected back to the volume and intersected with the cortical layers to obtain deep, middle and superficial portions of each region.

In addition to the neocortical ROIs, hippocampal subfields were defined bilaterally in the native anatomical space of each subject using the Hippocampal Segmentation Factory (HSF) toolbox (https://hsf.readthedocs.io/en/latest/) [68]. The resulting segmentation included the subiculum (SUB), Cornu Ammonis subfields (CA1, CA2, and CA3), and the dentate gyrus (DG).

#### 1.5.3 Single-trial Response Estimation

All analyses were conducted in the subject native space (0.8 mm isotropic resolution). We estimated trial-specific beta activation (expressed as percent signal change) using GLMSingle [65] for the visual (i.e., the three leading pictures) and the auditory (or omission thereof) portion of every trial, as well as the cued (i.e., delayed) motor response. All the analyses we report here are performed on the estimates of the response to the sound portion of each trial. For the functional connectivity analyses, the single-trial estimation model (GLMsingle) also incorporated data-driven nuisance regressors estimated with GLMdenoise together with ridge regularization [94] to improve the robustness of single-trial estimates while reducing the overlap between neighbouring haemodynamic responses and, consequently, autocorrelation within the beta series.

#### 1.5.4 Univariate fMRI Data Analysis

For each participant, we selected voxels whose overall activation was significant (omnibus test on the response to all auditory trials compared to baseline; p*<*0.05, uncorrected, t(287)*>*1.96). Note that voxel selection was based on overall task responsiveness and was therefore orthogonal to subsequent analyses, which are based on a differential contrast between predictable and unpredictable conditions. Within each layer of cortical ROIs, responses to valid and invalid trials were averaged separately across the selected voxels for each participant. Responses within corresponding laminar ROIs were averaged across hemispheres, as no specific hypotheses regarding hemispheric lateralisation were formulated. Individual subject estimates were used to test the cortical-depth *×* condition interaction using a non-parametric Friedman test to assess whether the effect of validity varied across layers.

#### 1.5.5 Neocortical ROI Selection

Neocortical regions of interest (ROIs) were restricted to a sub-network of areas that showed a significant layer-specific validity effect (corrected for multiple comparisons using FDR) in the preliminary univariate analysis (see Supplementary Table 1 and Supplementary Figure 5). In addition to the four cortical regions showing significant interaction effects (pSTG, pMTG, pars Orbitalis and pars Triangularis), we included TPOj (broadly covering the Temporal-Parietal-Occipital junction and the Supra Marginal Gyrus) on theoretical grounds because of its proposed involvement in multisensory integration and cross-modal predictive processing, and because it has been previously implicated in the processing of predictable auditory content in the same paradigm [62].

#### 1.5.6 Connectivity Analysis

The functional connectivity analyses were restricted to predictable valid trials and unpredictable (sound present) trials. In each participant dataset, we further restricted the analyses to voxels whose response to these trials was significant (omnibus test: all valid predictable and sound unpredictable trials vs zero, p*<*0.05, uncorrected). For each participant, ROI-specific beta series were obtained by averaging the single-trial estimates across the selected voxels of each cortical layer (deep, middle, and superficial) and hippocampal subfields (CA1-CA3, DG, and SUB). We therefore obtained, for both predictable and unpredictable conditions, beta-series matrices including hippocampal subfields and the three cortical layers of each neocortical ROI (Figure 1). Hippocampal subfields served as seed regions, whereas layer-specific cortical ROIs constituted the target regions of the functional connectivity analysis.

We estimated functional connectivity between hippocampal subfields and laminar cortical regions separately for the predictable and unpredictable conditions using beta series correlation [95] (Figure 1). To ensure an equal number of trials across predictable and unpredictable conditions, a bootstrap procedure (*n* = 1000) was used to subsample trials from the predictable valid condition to match the number of unpredictable (sound) trials. Beta series were z-scored within each region to remove differences in scale and mean activity level across regions. For each bootstrap sample, pairwise Pearson correlations were computed between hippocampal and cortical beta series and subsequently Fisher z-transformed. The resulting Fisher z-transformed connectivity matrices were averaged across bootstrap iterations to obtain a representative connectivity matrix for the predictable condition. Connectivity for the unpredictable condition was estimated analogously from all available unpredictable trials. Because no specific hypotheses regarding hemispheric lateralisation were formulated, connectivity estimates involving homologous cortical areas were averaged across hemispheres in both conditions.

To assess the modulation of functional connectivity induced by predictability, we computed, for each participant, the difference between Fisher z-transformed connectivity estimates in the two conditions: (Δ*z*_BSC_ = *z*_predictable_ *− z*_unpredictable_).

Group-level statistical inference was based on the within-subject difference and tested edgewise using a non-parametric sign-flipping permutation test. Resulting p-values were corrected for multiple comparisons using the Benjamini-Hochberg false discovery rate procedure (*q*_FDR_¡0.05) [96].

We additionally analysed region-level connectivity estimates, obtained without sub-dividing cortical ROIs by depth. Region-level beta-series correlations were estimated using the same procedure described above and Fisher *z*-transformed before statistical analysis.

## Acknowledgements

We would like to thank Giancarlo Valente for his advice on permutation testing and statistical tests and Jorie van Haren for feedback on the preprocessing pipeline. The authors thank all members of the lab for their support.

## Author contributions

Conceptualization, F.D.M., S.S., F.E., N.V.; methodology, F.D.M., S.S., M.E.; investigation, S.S., M.E., F.D.M.; writing – original draft, S.S., F.D.M.; writing – review & editing, M.E., R.A., L.M., F.E., N.V., C.S.; funding acquisition, F.D.M.; supervision, F.D.M., N.V.

## Materials & Correspondence

Correspondence and material requests should be addressed to m

## Data availability

All data are available at: https://openneuro.org/datasets/ds007109

## Code availability

The processing pipeline and custom analysis code used in this work are available at: https://github.com/mesoScopic-Computational-AuditioN-lab/ResearchProjects/ tree/main/finalized/AssociativeLearning.

## Supplementary Materials

### Cortical depth-dependent effect of validity

**Table 1:**
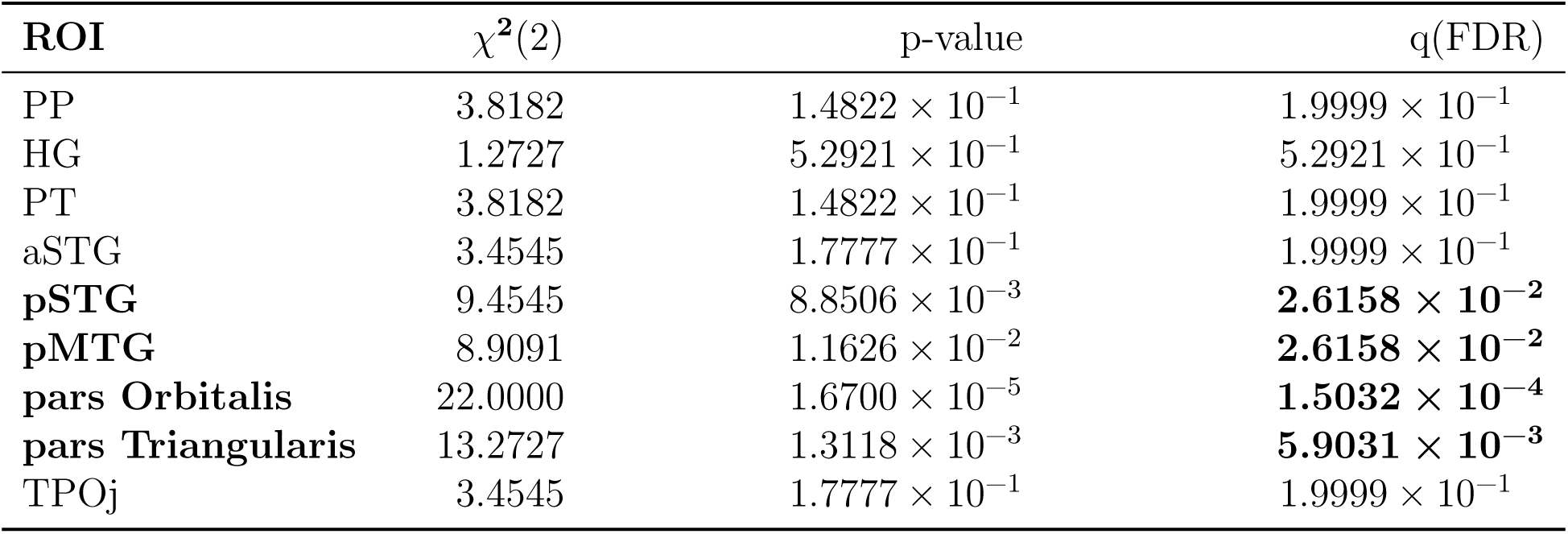
Analysis of the interaction between cortical layers and validity (valid vs. invalid trials - Friedman test - degrees of freedom reported in parentheses, correction for multiple comparison was preformed using False Discovery Rate)

### Cortical depth-dependent beta series correlation

**Table 2:**
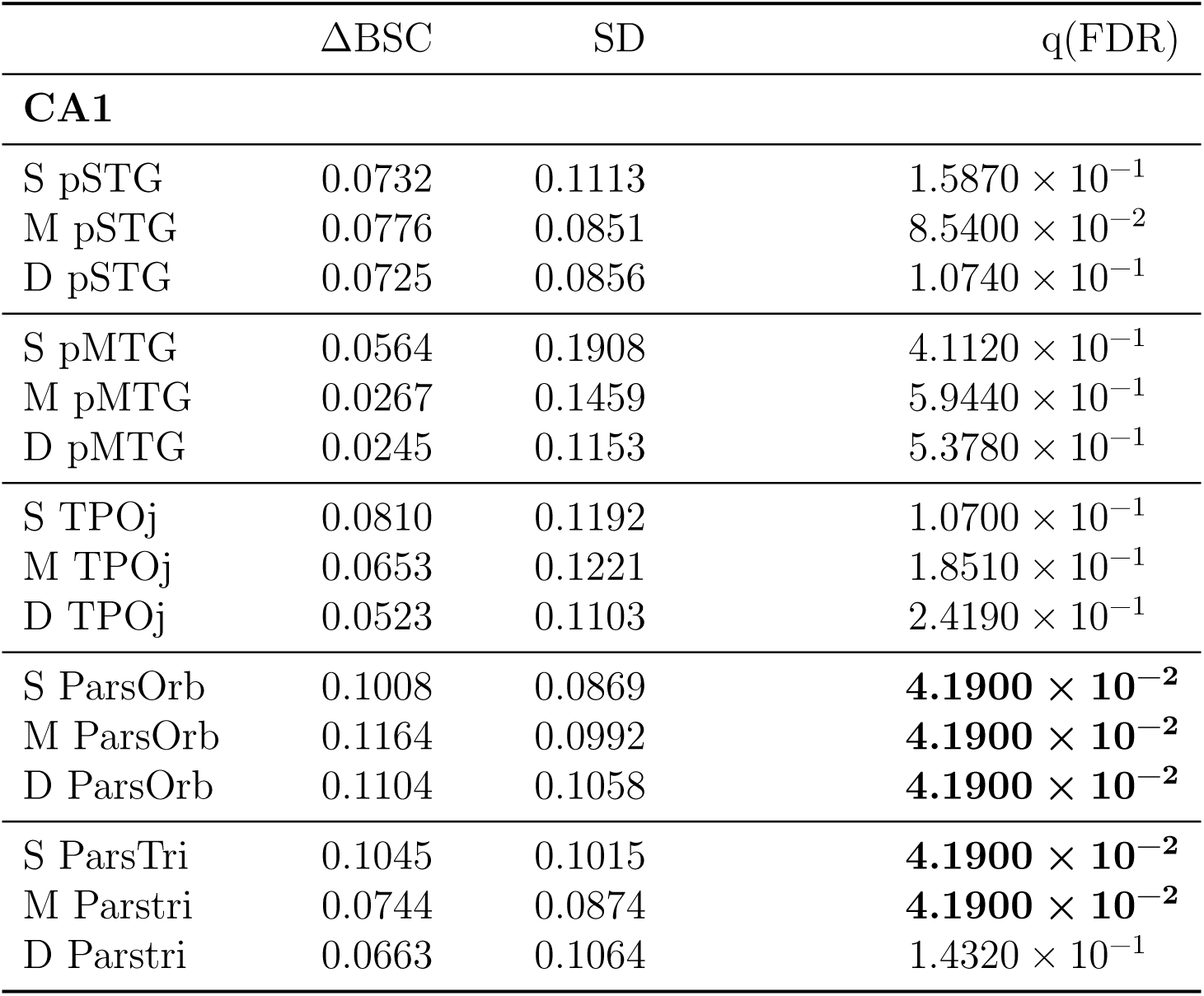
Mean difference (predictable - unpredictable) in beta series correlation (ΔBSC), its standard deviation across participants, and the corresponding FDR-corrected *q*-value are reported for the connectivity between CA1 and layers (deep [D], middle [M], and superficial [S]) of cortical regions.

**Table 3:**
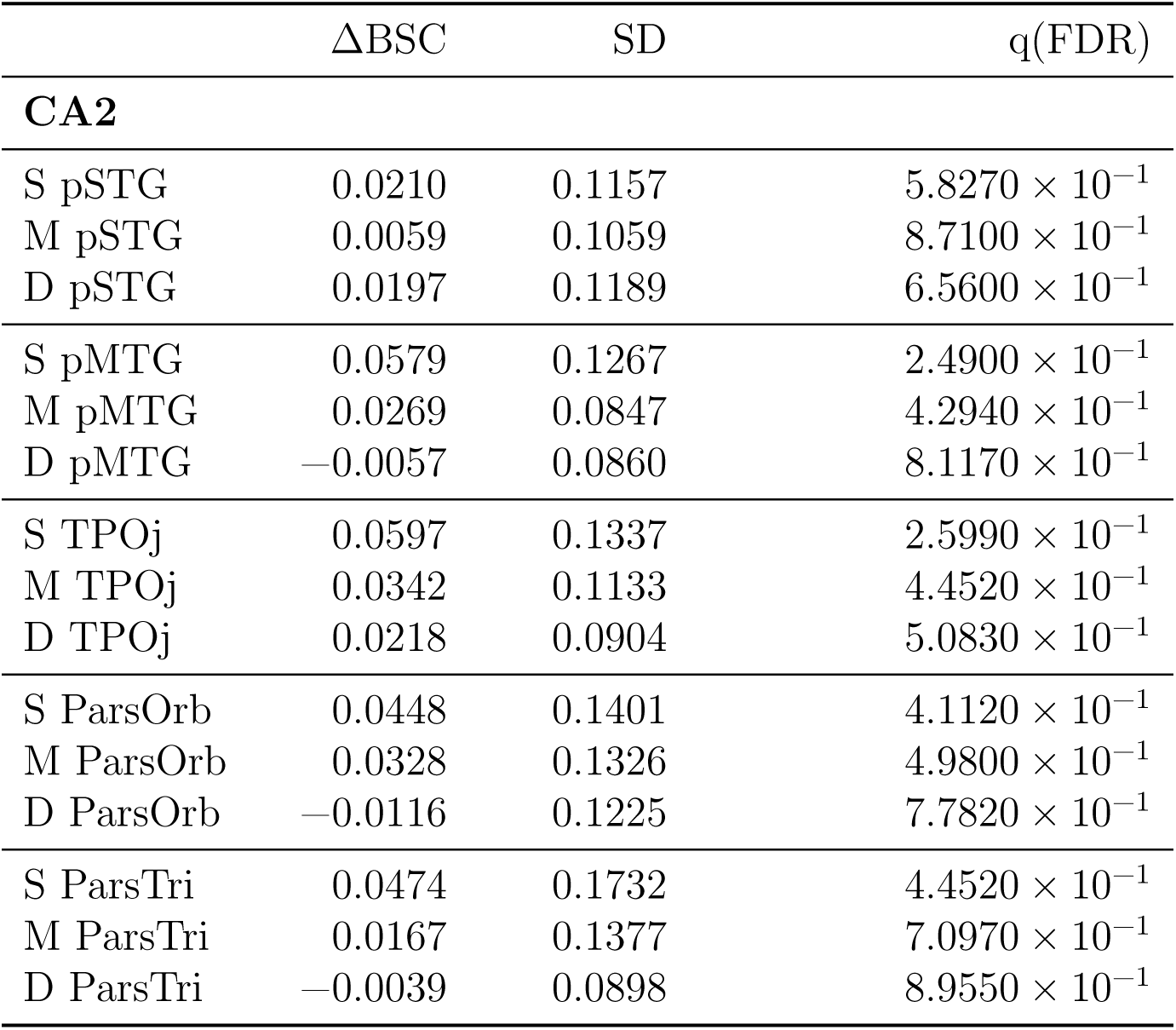
Mean difference (predictable - unpredictable) in beta series correlation (ΔBSC), its standard deviation across participants, and the corresponding FDR-corrected *q*-value are reported for the connectivity between CA2 and layers (deep [D], middle [M], and superficial [S]) of cortical regions.

**Table 4:**
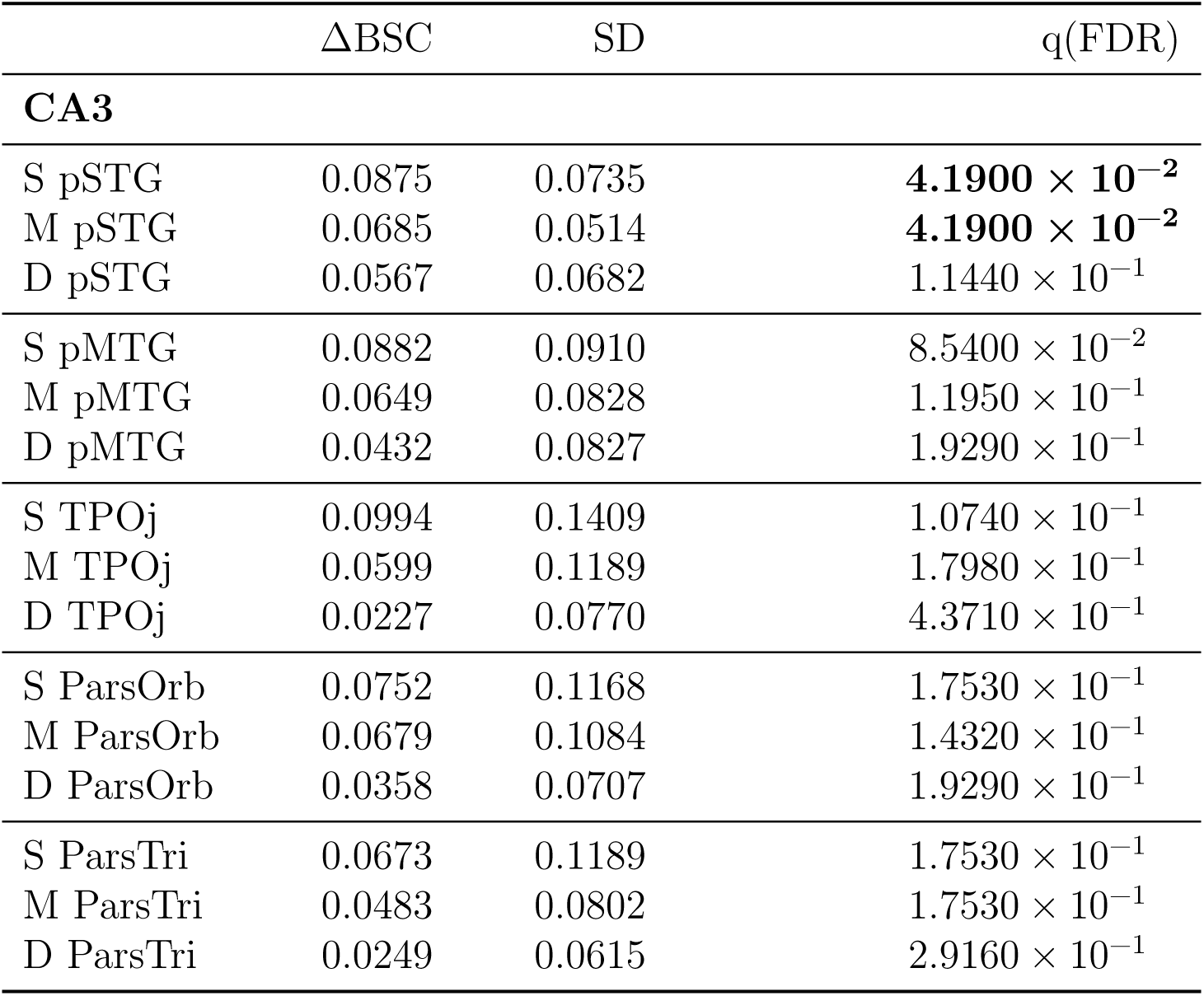
Mean difference (predictable - unpredictable) in beta series correlation (ΔBSC), its standard deviation across participants, and the corresponding FDR-corrected *q*-value are reported for the connectivity between CA3 and layers (deep [D], middle [M], and superficial [S]) of cortical regions.

**Table 5:**
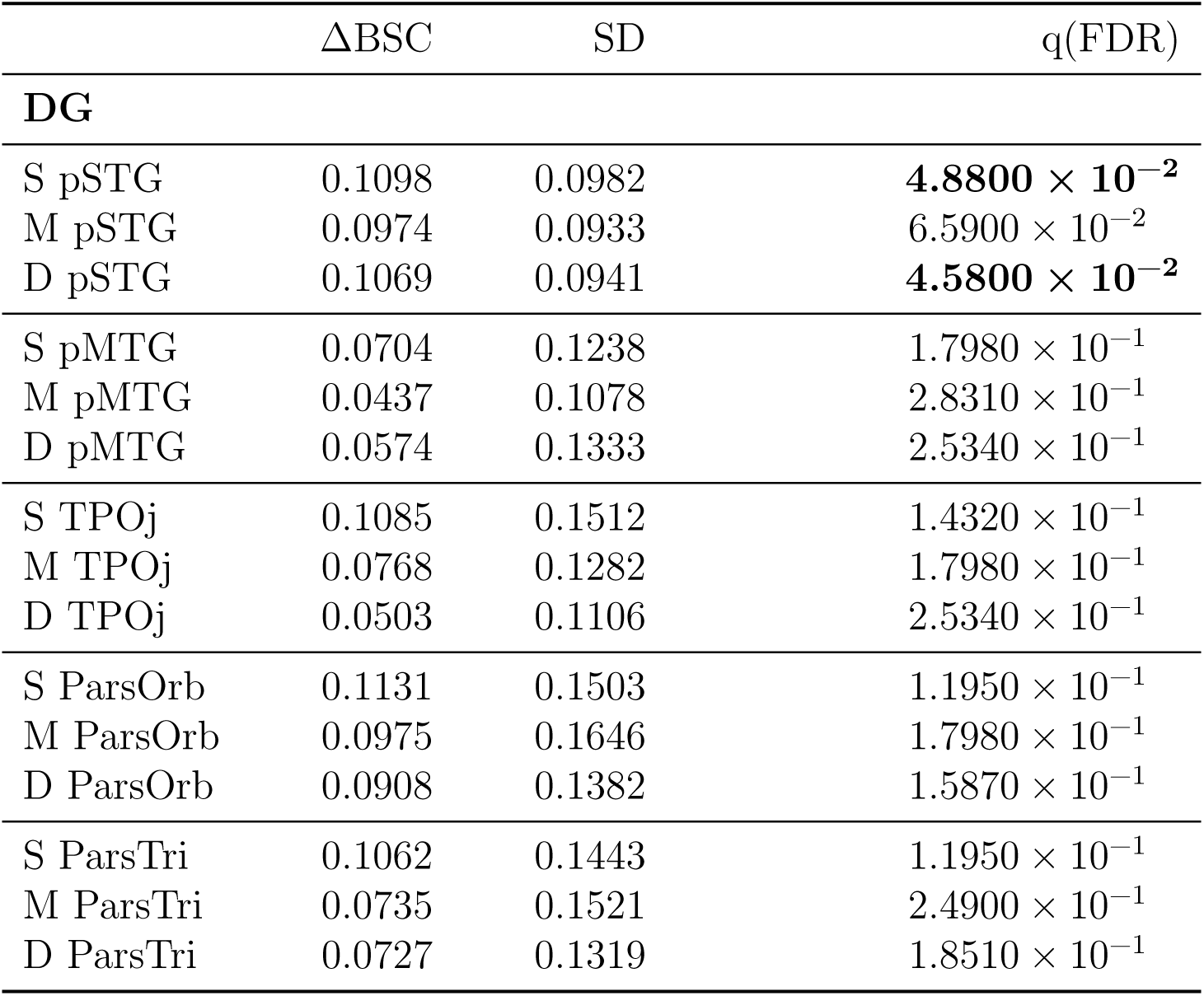
Mean difference (predictable - unpredictable) in beta series correlation (ΔBSC), its standard deviation across participants, and the corresponding FDR-corrected *q*-value are reported for the connectivity between DG and layers (deep [D], middle [M], and superficial [S]) of cortical regions.

**Table 6:**
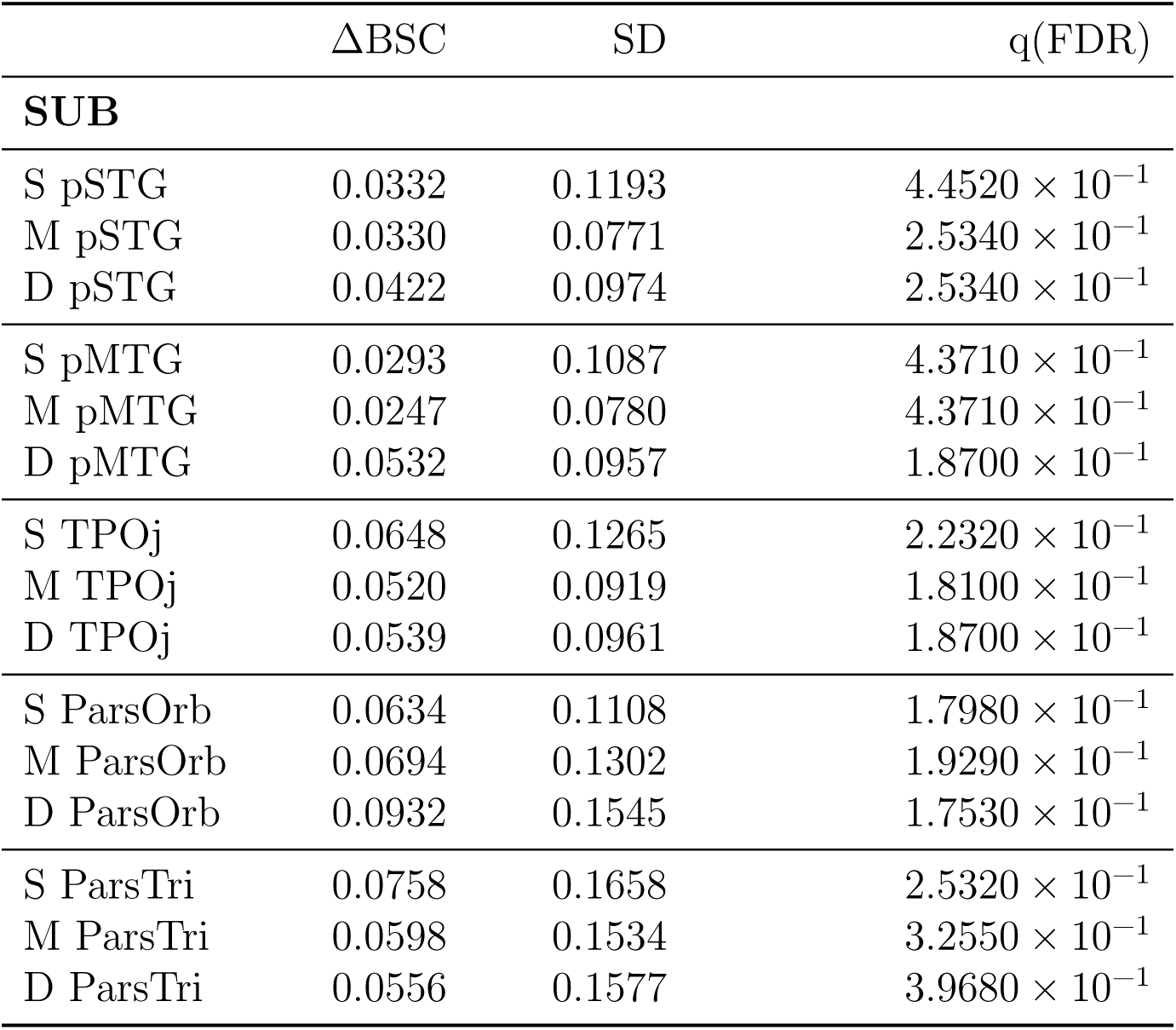
Difference (Mean difference (predictable - unpredictable) in beta series correlation (ΔBSC), its standard deviation across participants, and the corresponding FDR-corrected *q*-value are reported for the connectivity between the subiculum and layers (deep [D], middle [M], and superficial [S]) of cortical regions.

**Figure 4:**
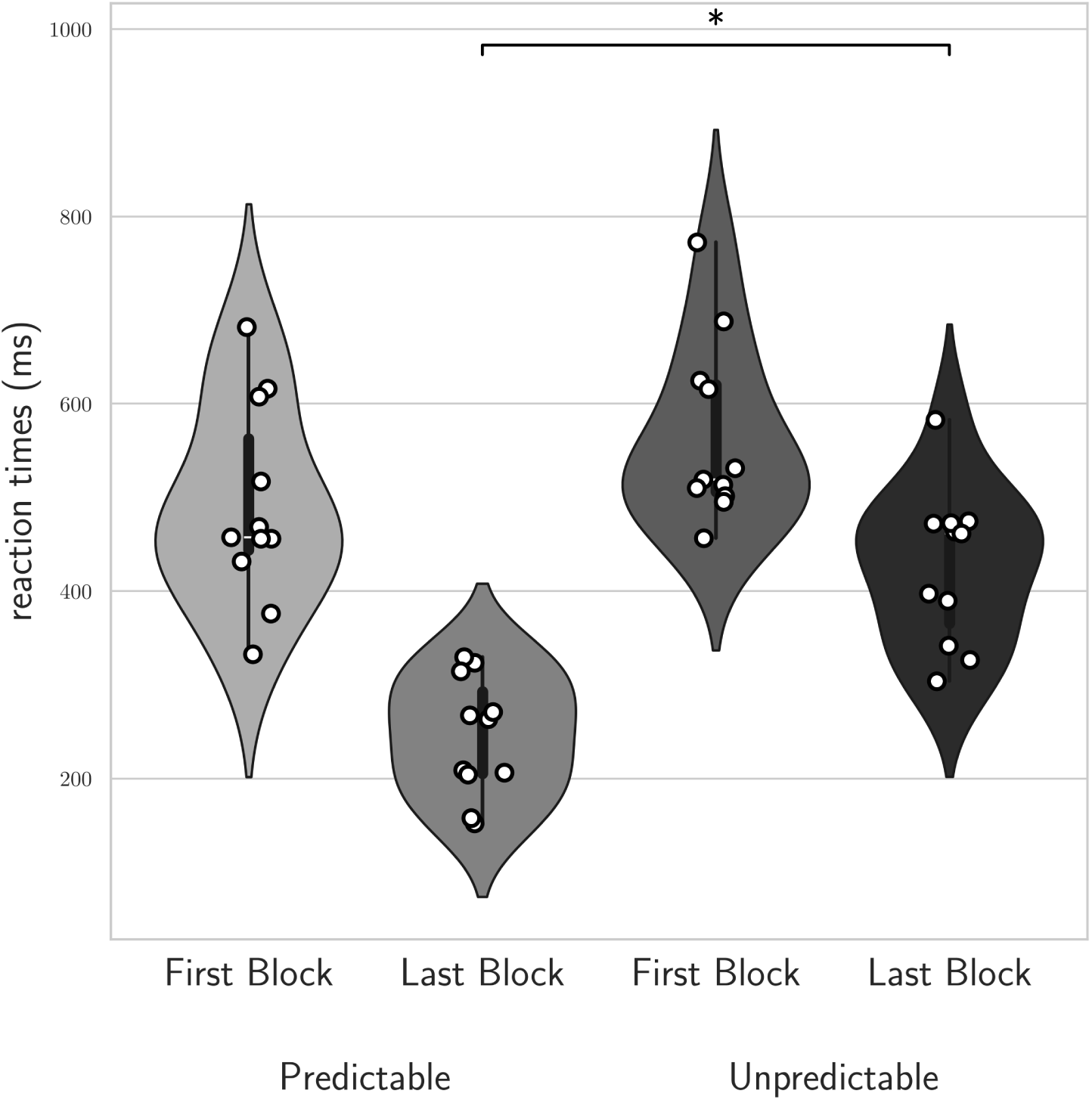
Behavioral performance during training. Mean reaction times for predictable and unpredictable trials in the first and last training blocks across all participants (N = 11). Violin plots indicate group distributions, and white circles show individual participants’ mean reaction times averaged across trials. The asterisk denotes a significant interaction between training block (first vs. last) and context (predictable vs. unpredictable).

**Supplementary Figure 5:**
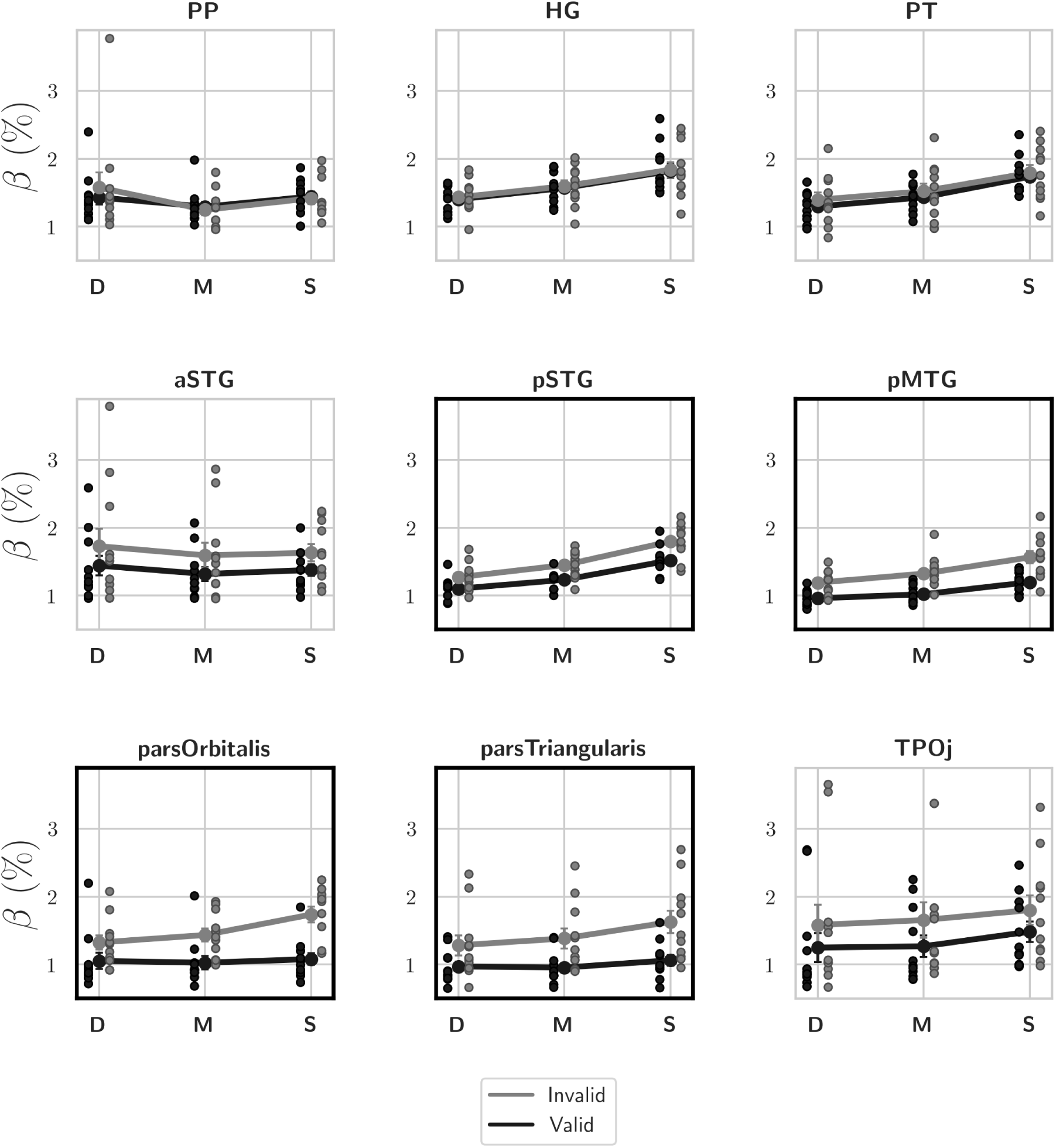
Effect of validity cortical layers and ROIs. Layer-dependent responses were measured for valid (black) and invalid (gray) trials across cortical ROIs: planum polare (PP), Heschl’s gyrus (HG), planum temporale (PT), anterior and posterior superior temporal gyrus (aSTG, pSTG), posterior middle temporal gyrus (pMTG), pars triangularis, pars orbitalis, and temporo-parietal junction (TPOj). Layers are deep [D], middle [M], and superficial [S]; plots show the average percent signal change across participants for each condition (error bars = standard error; dots represent individual data). Significant condition*×*layer interactions are highlighted with a bold outline.

**Supplementary Figure 6:**
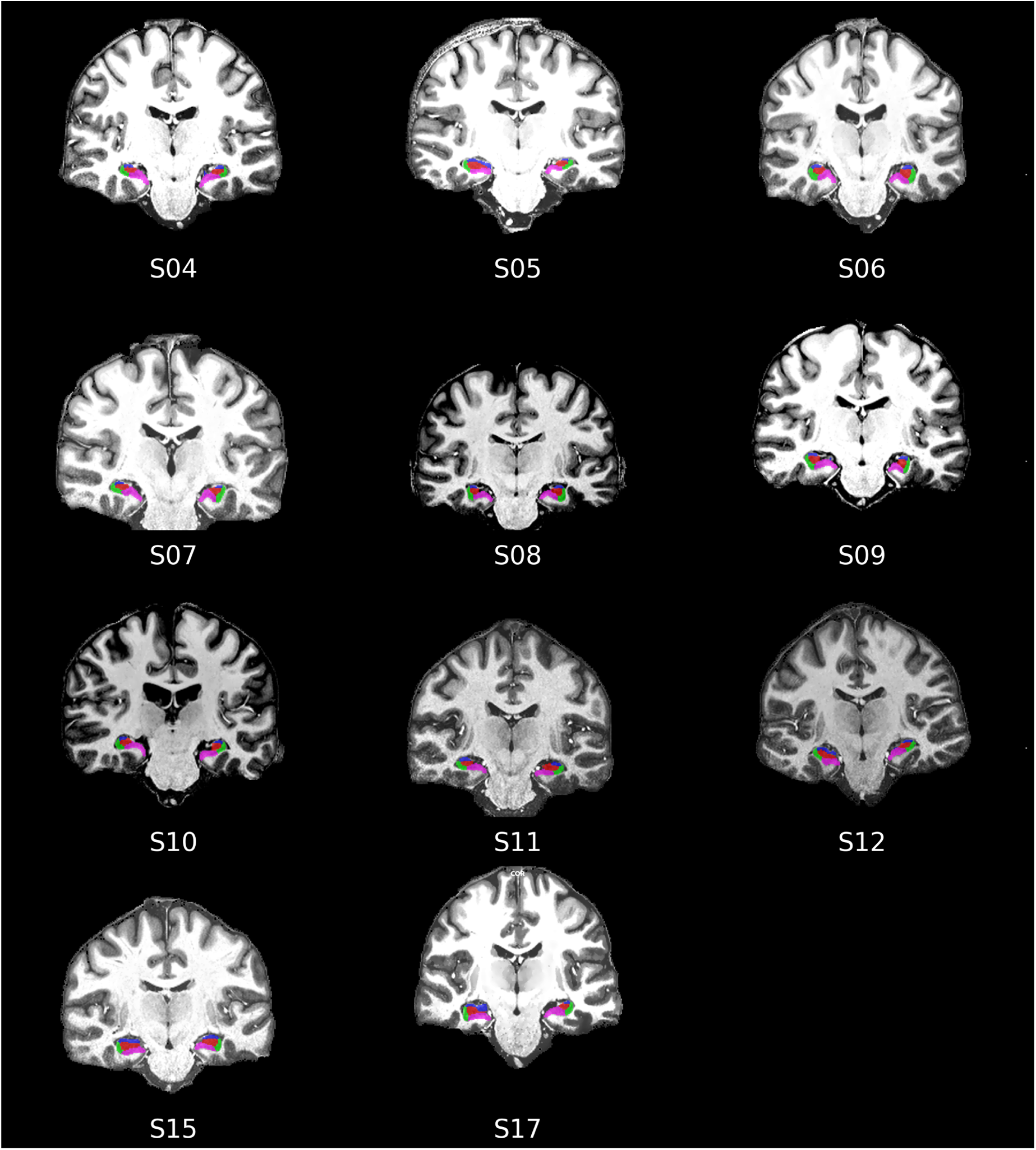
Individual hippocampal ROIs. We used the Hippocampal Segmentation Factory toolbox [68] to define hippocampal subfields (cornu ammonis [CA1, CA2 CA3]), dentate gyrus (DG) and subiculum (Sub) in the native anatomical space of each participant.

**Supplementary Figure 7:**
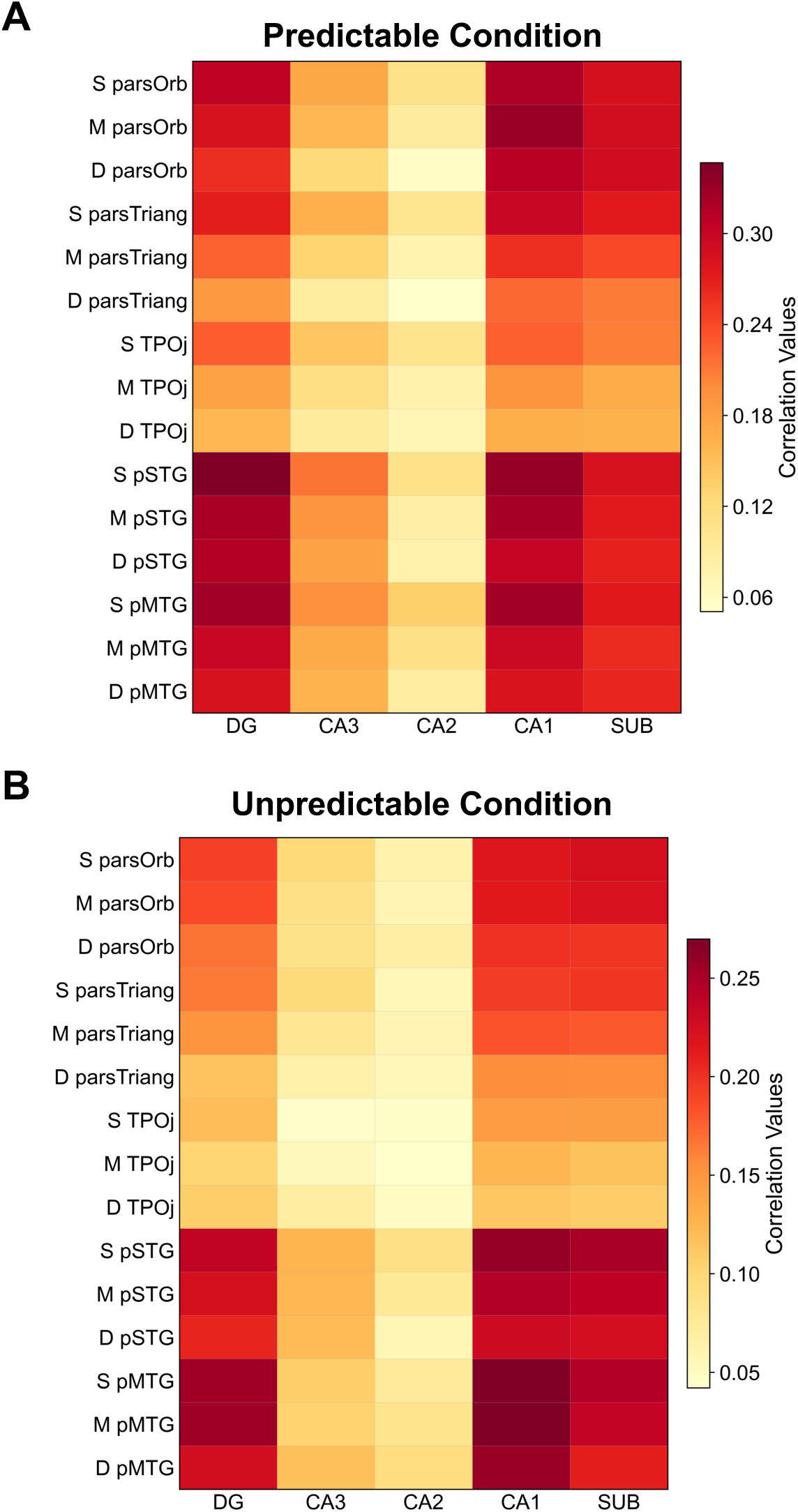
Laminar-level beta series correlation in predictable and unpredictable conditions. Unthresholded heatmaps of beta series correlation values averaged across subjects for the predictable *(A)* and unpredictable *(B))* conditions. Rows: cortical ROIs across superficial (S), middle (M), and deep (D) layers; columns: hippocampal subfields.

**Supplementary Figure 8:**
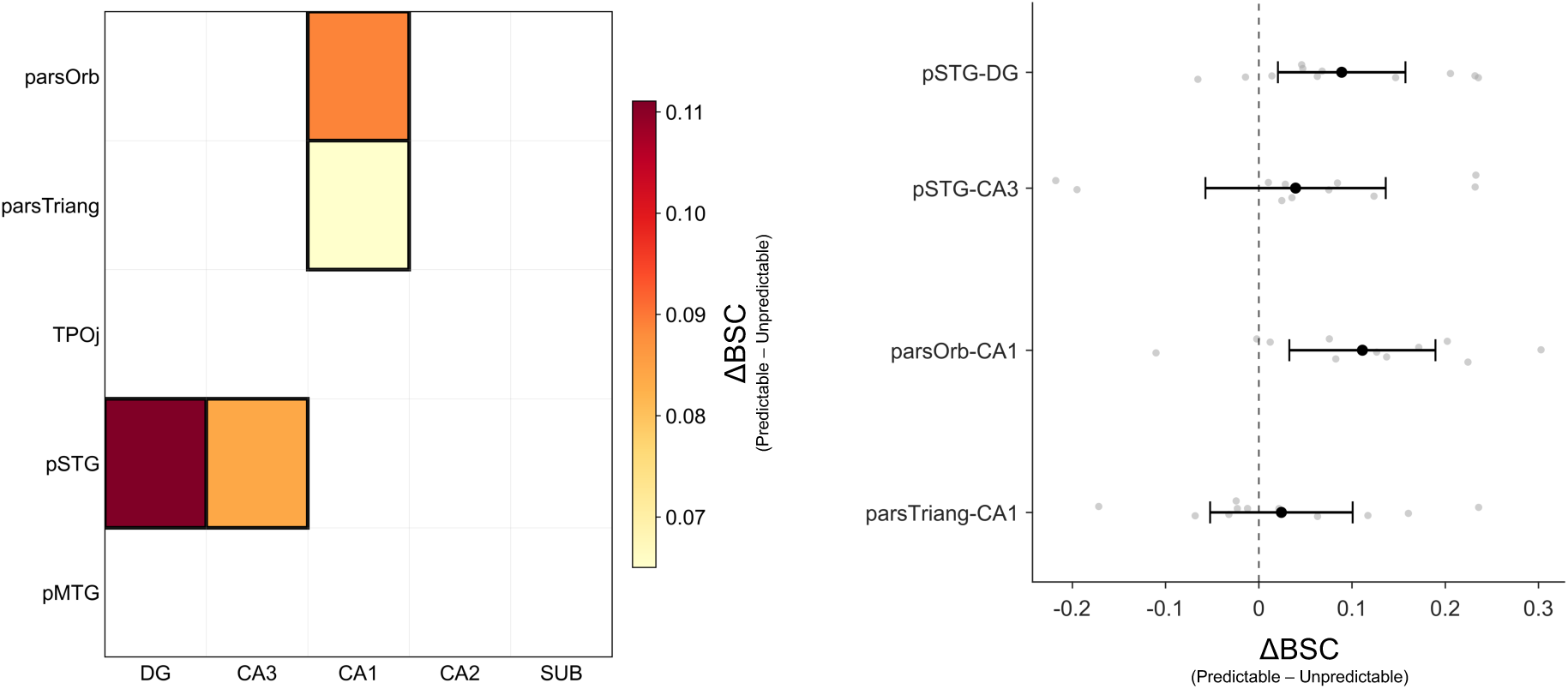
Beta-series connectivity at the level of whole cortical ROIs. (*A*) Heatmap of group-level significant predictability effects on regional (whole-ROI) BSC connectivity after collapsing across cortical depth. Cells show the mean connectivity difference (predictable - unpredictable) for ROI pairs surviving FDR correction (*q*_FDR_ *<* 0.05). Cortical ROIs (pSTG, pMTG, TPOj, pars orbitalis, pars triangularis) and hippocampal subfields (CA1, CA3, DG, SUB) are shown on the two axes. (*B*) Uncertainty of the predictability-related effects. Individual participant values are shown for the difference in beta-series correlation between predictable and unpredictable conditions (ΔBSC = BSC_predictable_ *−* BSC_unpredictable_). Black dots indicate group means and horizontal error bars the corresponding 95% confidence intervals. Only edges surviving non-parametric sign-flipping permutation testing with FDR correction (*q*_FDR_ *<* 0.05) are shown.

**Supplementary Figure 9:**
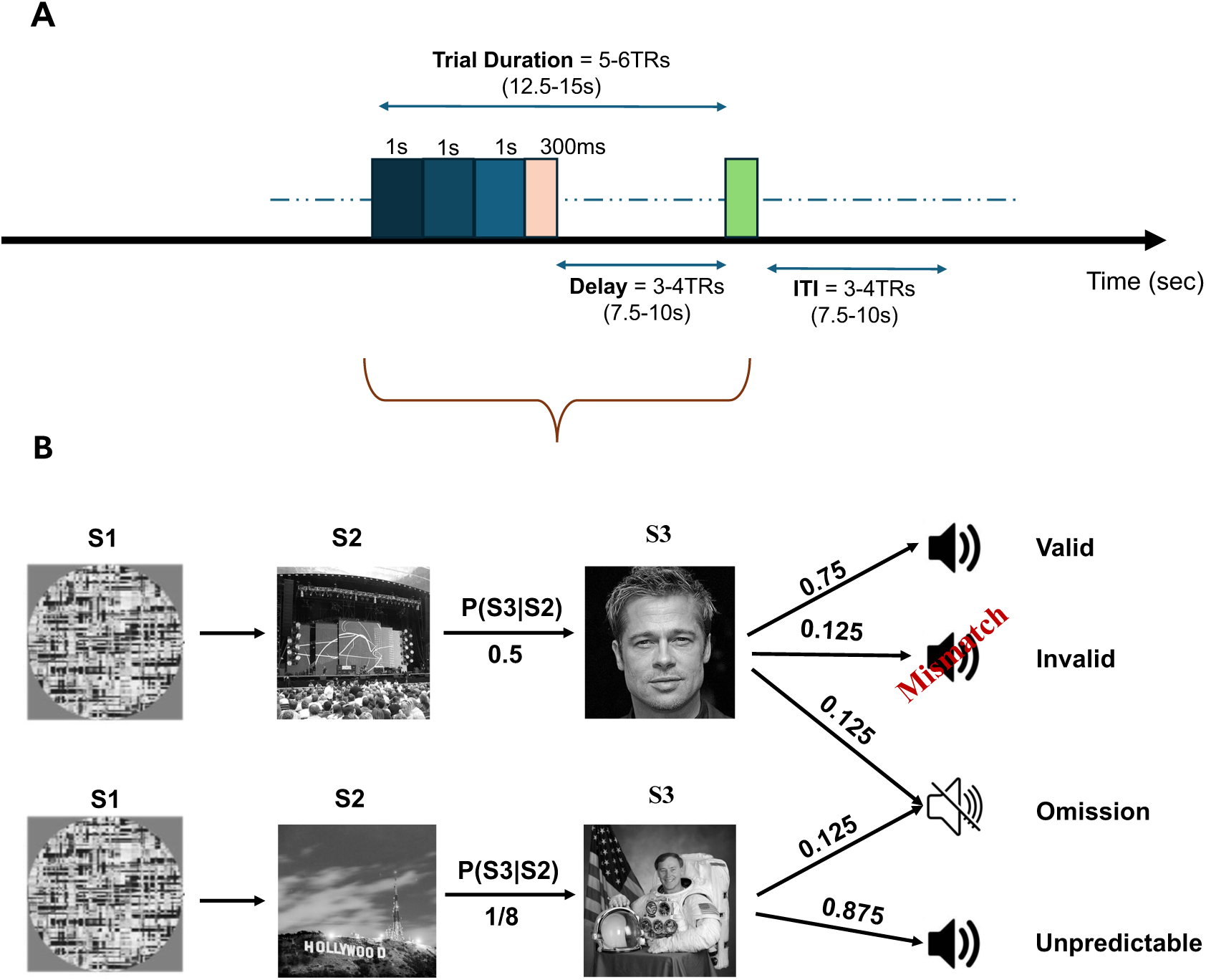
Experimental design. *(A)* During fMRI acquisition participants listened, in each trial, to an auditory syllable (pink bar, 300 ms) following visual pictures (three pictures presented for one second each). Every trial ended with a cued participant response (green bar) and trials had a random intertrial interval of 3-4 TRs. *(B)* In predictable trials, the auditory syllable could be predicted from the preceding on the basis of a preceding behavioral training. We presented the expected (valid) syllable in 75% of trials, the remaining trials were equally divided between invalid and omission. In unpredictable trials, visual stimuli did not predict the syllable, and omission trials were also included (12.5%).

