## supplementary materials for "Learned Audiovisual Predictability Modulates Depth-Resolved Hippocampal–Cortical Functional Coupling"

October 16, 2026

**Cortical depth-dependent effect of validity**

| **ROI** | $\chi^2(2)$ | p-value | q(FDR) |
|---|---|---|---|
| PP | 3.8182 | $1.4822 \times 10^{-1}$ | $1.9999 \times 10^{-1}$ |
| HG | 1.2727 | $5.2921 \times 10^{-1}$ | $5.2921 \times 10^{-1}$ |
| PT | 3.8182 | $1.4822 \times 10^{-1}$ | $1.9999 \times 10^{-1}$ |
| aSTG | 3.4545 | $1.7777 \times 10^{-1}$ | $1.9999 \times 10^{-1}$ |
| **pSTG** | 9.4545 | $8.8506 \times 10^{-3}$ | $\mathbf{2.6158 \times 10^{-2}}$ |
| **pMTG** | 8.9091 | $1.1626 \times 10^{-2}$ | $\mathbf{2.6158 \times 10^{-2}}$ |
| **pars Orbitalis** | 22.0000 | $1.6700 \times 10^{-5}$ | $\mathbf{1.5032 \times 10^{-4}}$ |
| **pars Triangularis** | 13.2727 | $1.3118 \times 10^{-3}$ | $\mathbf{5.9031 \times 10^{-3}}$ |
| TPOj | 3.4545 | $1.7777 \times 10^{-1}$ | $1.9999 \times 10^{-1}$ |

Table 1: Analysis of the interaction between cortical layers and validity (valid vs. invalid trials - Friedman test - degrees of freedom reported in parentheses, correction for multiple comparison was preformed using False Discovery Rate)

**Cortical depth-dependent beta series correlation**

| | ΔBSC | SD | q(FDR) |
|---|---|---|---|
| **CA1** | | | |
| S pSTG | 0.0732 | 0.1113 | $1.5870 \times 10^{-1}$ |
| M pSTG | 0.0776 | 0.0851 | $8.5400 \times 10^{-2}$ |
| D pSTG | 0.0725 | 0.0856 | $1.0740 \times 10^{-1}$ |
| S pMTG | 0.0564 | 0.1908 | $4.1120 \times 10^{-1}$ |
| M pMTG | 0.0267 | 0.1459 | $5.9440 \times 10^{-1}$ |
| D pMTG | 0.0245 | 0.1153 | $5.3780 \times 10^{-1}$ |
| S TPOj | 0.0810 | 0.1192 | $1.0700 \times 10^{-1}$ |
| M TPOj | 0.0653 | 0.1221 | $1.8510 \times 10^{-1}$ |
| D TPOj | 0.0523 | 0.1103 | $2.4190 \times 10^{-1}$ |
| S ParsOrb | 0.1008 | 0.0869 | $\mathbf{4.1900 \times 10^{-2}}$ |
| M ParsOrb | 0.1164 | 0.0992 | $\mathbf{4.1900 \times 10^{-2}}$ |
| D ParsOrb | 0.1104 | 0.1058 | $\mathbf{4.1900 \times 10^{-2}}$ |
| S ParsTri | 0.1045 | 0.1015 | $\mathbf{4.1900 \times 10^{-2}}$ |
| M Parstri | 0.0744 | 0.0874 | $\mathbf{4.1900 \times 10^{-2}}$ |
| D Parstri | 0.0663 | 0.1064 | $1.4320 \times 10^{-1}$ |

Table 2: Mean difference (predictable - unpredictable) in beta series correlation (ΔBSC), its standard deviation across participants, and the corresponding FDR-corrected $q$-value are reported for the connectivity between CA1 and layers (deep [D], middle [M], and superficial [S]) of cortical regions.

| | ΔBSC | SD | q(FDR) |
|---|---|---|---|
| **CA2** | | | |
| S pSTG | 0.0210 | 0.1157 | $5.8270 \times 10^{-1}$ |
| M pSTG | 0.0059 | 0.1059 | $8.7100 \times 10^{-1}$ |
| D pSTG | 0.0197 | 0.1189 | $6.5600 \times 10^{-1}$ |
| S pMTG | 0.0579 | 0.1267 | $2.4900 \times 10^{-1}$ |
| M pMTG | 0.0269 | 0.0847 | $4.2940 \times 10^{-1}$ |
| D pMTG | −0.0057 | 0.0860 | $8.1170 \times 10^{-1}$ |
| S TPOj | 0.0597 | 0.1337 | $2.5990 \times 10^{-1}$ |
| M TPOj | 0.0342 | 0.1133 | $4.4520 \times 10^{-1}$ |
| D TPOj | 0.0218 | 0.0904 | $5.0830 \times 10^{-1}$ |
| S ParsOrb | 0.0448 | 0.1401 | $4.1120 \times 10^{-1}$ |
| M ParsOrb | 0.0328 | 0.1326 | $4.9800 \times 10^{-1}$ |
| D ParsOrb | −0.0116 | 0.1225 | $7.7820 \times 10^{-1}$ |
| S ParsTri | 0.0474 | 0.1732 | $4.4520 \times 10^{-1}$ |
| M ParsTri | 0.0167 | 0.1377 | $7.0970 \times 10^{-1}$ |
| D ParsTri | −0.0039 | 0.0898 | $8.9550 \times 10^{-1}$ |

Table 3: Mean difference (predictable - unpredictable) in beta series correlation (ΔBSC), its standard deviation across participants, and the corresponding FDR-corrected *q*-value are reported for the connectivity between CA2 and layers (deep [D], middle [M], and superficial [S]) of cortical regions.

| | ΔBSC | SD | q(FDR) |
|---|---|---|---|
| **CA3** | | | |
| S pSTG | 0.0875 | 0.0735 | $\mathbf{4.1900 \times 10^{-2}}$ |
| M pSTG | 0.0685 | 0.0514 | $\mathbf{4.1900 \times 10^{-2}}$ |
| D pSTG | 0.0567 | 0.0682 | $1.1440 \times 10^{-1}$ |
| S pMTG | 0.0882 | 0.0910 | $8.5400 \times 10^{-2}$ |
| M pMTG | 0.0649 | 0.0828 | $1.1950 \times 10^{-1}$ |
| D pMTG | 0.0432 | 0.0827 | $1.9290 \times 10^{-1}$ |
| S TPOj | 0.0994 | 0.1409 | $1.0740 \times 10^{-1}$ |
| M TPOj | 0.0599 | 0.1189 | $1.7980 \times 10^{-1}$ |
| D TPOj | 0.0227 | 0.0770 | $4.3710 \times 10^{-1}$ |
| S ParsOrb | 0.0752 | 0.1168 | $1.7530 \times 10^{-1}$ |
| M ParsOrb | 0.0679 | 0.1084 | $1.4320 \times 10^{-1}$ |
| D ParsOrb | 0.0358 | 0.0707 | $1.9290 \times 10^{-1}$ |
| S ParsTri | 0.0673 | 0.1189 | $1.7530 \times 10^{-1}$ |
| M ParsTri | 0.0483 | 0.0802 | $1.7530 \times 10^{-1}$ |
| D ParsTri | 0.0249 | 0.0615 | $2.9160 \times 10^{-1}$ |

Table 4: Mean difference (predictable - unpredictable) in beta series correlation (ΔBSC), its standard deviation across participants, and the corresponding FDR-corrected *q*-value are reported for the connectivity between CA3 and layers (deep [D], middle [M], and superficial [S]) of cortical regions.

| | ΔBSC | SD | q(FDR) |
|---|---|---|---|
| **DG** | | | |
| S pSTG | 0.1098 | 0.0982 | $\mathbf{4.8800 \times 10^{-2}}$ |
| M pSTG | 0.0974 | 0.0933 | $6.5900 \times 10^{-2}$ |
| D pSTG | 0.1069 | 0.0941 | $\mathbf{4.5800 \times 10^{-2}}$ |
| S pMTG | 0.0704 | 0.1238 | $1.7980 \times 10^{-1}$ |
| M pMTG | 0.0437 | 0.1078 | $2.8310 \times 10^{-1}$ |
| D pMTG | 0.0574 | 0.1333 | $2.5340 \times 10^{-1}$ |
| S TPOj | 0.1085 | 0.1512 | $1.4320 \times 10^{-1}$ |
| M TPOj | 0.0768 | 0.1282 | $1.7980 \times 10^{-1}$ |
| D TPOj | 0.0503 | 0.1106 | $2.5340 \times 10^{-1}$ |
| S ParsOrb | 0.1131 | 0.1503 | $1.1950 \times 10^{-1}$ |
| M ParsOrb | 0.0975 | 0.1646 | $1.7980 \times 10^{-1}$ |
| D ParsOrb | 0.0908 | 0.1382 | $1.5870 \times 10^{-1}$ |
| S ParsTri | 0.1062 | 0.1443 | $1.1950 \times 10^{-1}$ |
| M ParsTri | 0.0735 | 0.1521 | $2.4900 \times 10^{-1}$ |
| D ParsTri | 0.0727 | 0.1319 | $1.8510 \times 10^{-1}$ |

Table 5: Mean difference (predictable - unpredictable) in beta series correlation (ΔBSC), its standard deviation across participants, and the corresponding FDR-corrected $q$-value are reported for the connectivity between DG and layers (deep [D], middle [M], and superficial [S]) of cortical regions.

| | ΔBSC | SD | q(FDR) |
|---|---|---|---|
| **SUB** | | | |
| S pSTG | 0.0332 | 0.1193 | $4.4520 \times 10^{-1}$ |
| M pSTG | 0.0330 | 0.0771 | $2.5340 \times 10^{-1}$ |
| D pSTG | 0.0422 | 0.0974 | $2.5340 \times 10^{-1}$ |
| S pMTG | 0.0293 | 0.1087 | $4.3710 \times 10^{-1}$ |
| M pMTG | 0.0247 | 0.0780 | $4.3710 \times 10^{-1}$ |
| D pMTG | 0.0532 | 0.0957 | $1.8700 \times 10^{-1}$ |
| S TPOj | 0.0648 | 0.1265 | $2.2320 \times 10^{-1}$ |
| M TPOj | 0.0520 | 0.0919 | $1.8100 \times 10^{-1}$ |
| D TPOj | 0.0539 | 0.0961 | $1.8700 \times 10^{-1}$ |
| S ParsOrb | 0.0634 | 0.1108 | $1.7980 \times 10^{-1}$ |
| M ParsOrb | 0.0694 | 0.1302 | $1.9290 \times 10^{-1}$ |
| D ParsOrb | 0.0932 | 0.1545 | $1.7530 \times 10^{-1}$ |
| S ParsTri | 0.0758 | 0.1658 | $2.5320 \times 10^{-1}$ |
| M ParsTri | 0.0598 | 0.1534 | $3.2550 \times 10^{-1}$ |
| D ParsTri | 0.0556 | 0.1577 | $3.9680 \times 10^{-1}$ |

Table 6: Difference (Mean difference (predictable - unpredictable) in beta series correlation (ΔBSC), its standard deviation across participants, and the corresponding FDR-corrected $q$-value are reported for the connectivity between the subiculum and layers (deep [D], middle [M], and superficial [S]) of cortical regions.

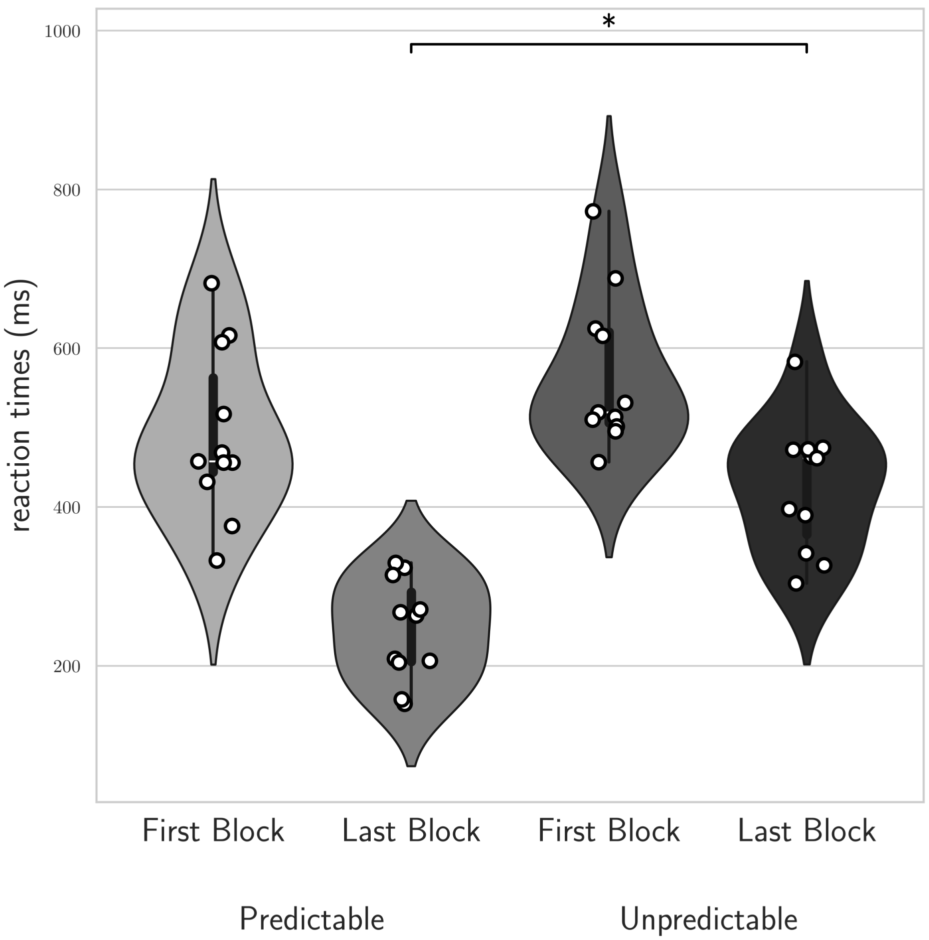


Figure 1: **Behavioral performance during training.** Mean reaction times for predictable and unpredictable trials in the first and last training blocks across all participants (N = 11). Violin plots indicate group distributions, and white circles show individual participants' mean reaction times averaged across trials. The asterisk denotes a significant interaction between training block (first vs. last) and context (predictable vs. unpredictable).

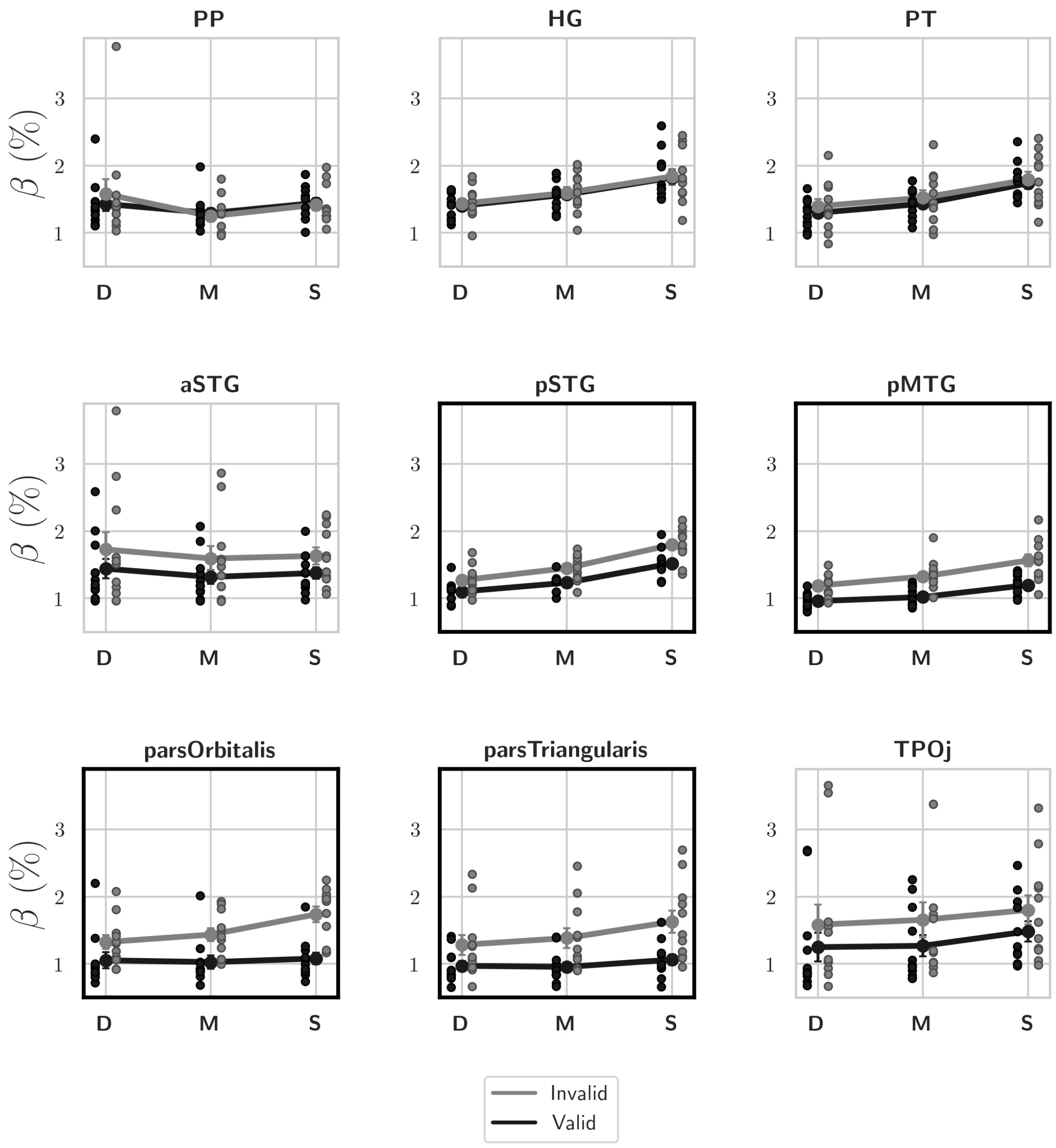


**Supplementary Figure 2: Effect of validity cortical layers and ROIs.** Layer-dependent responses were measured for valid (black) and invalid (gray) trials across cortical ROIs: planum polare (PP), Heschl's gyrus (HG), planum temporale (PT), anterior and posterior superior temporal gyrus (aSTG, pSTG), posterior middle temporal gyrus (pMTG), pars triangularis, pars orbitalis, and temporo-parietal junction (TPOj). Layers are deep [D], middle [M], and superficial [S]; plots show the average percent signal change across participants for each condition (error bars = standard error; dots represent individual data). Significant condition×layer interactions are highlighted with a bold outline.

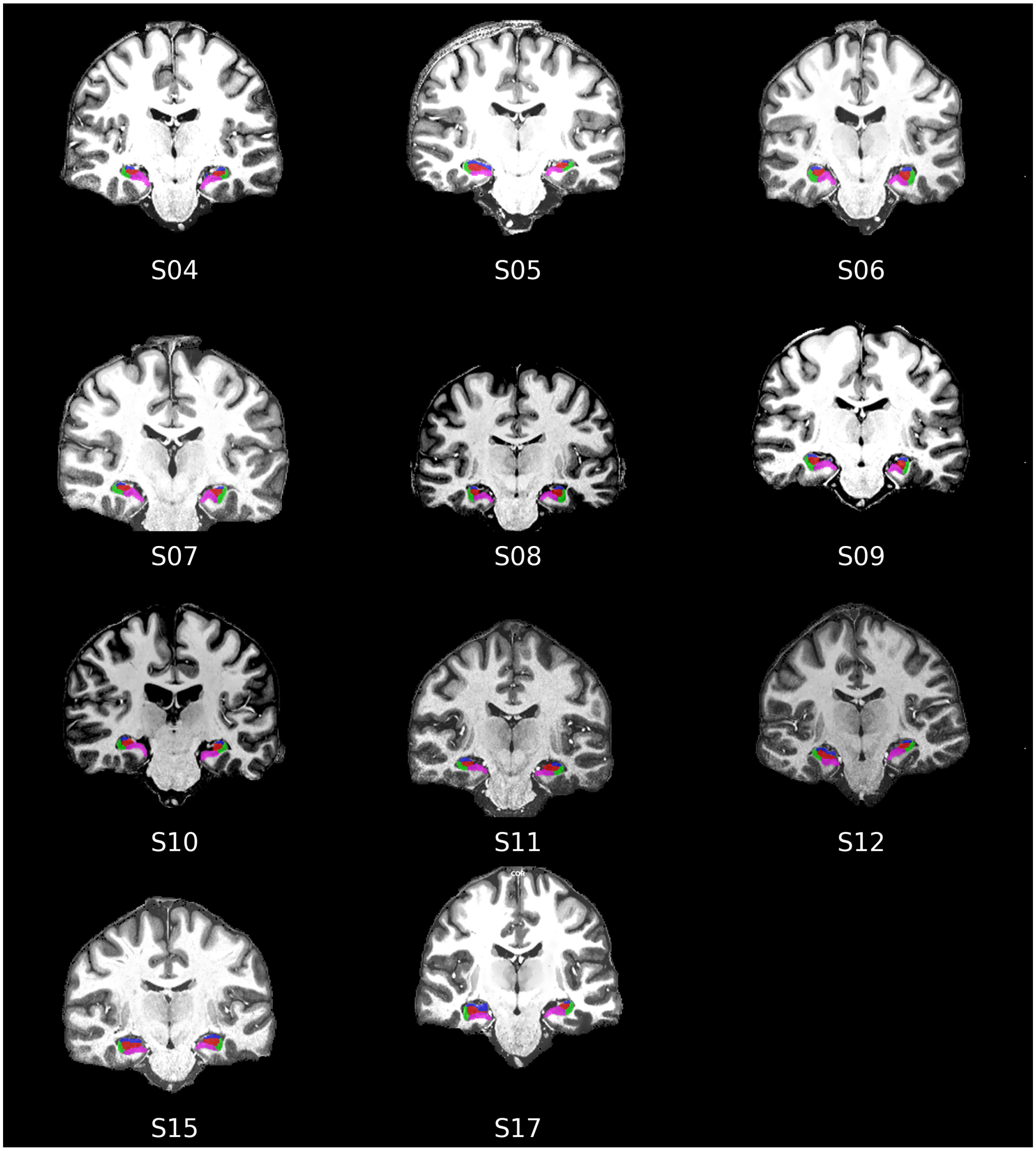


**Supplementary Figure 3: Individual hippocampal ROIs.** We used the Hippocampal Segmentation Factory toolbox to define hippocampal subfields (cornu ammonis [CA1, CA2 CA3]), dentate gyrus (DG) and subiculum (Sub) in the native anatomical space of each participant.

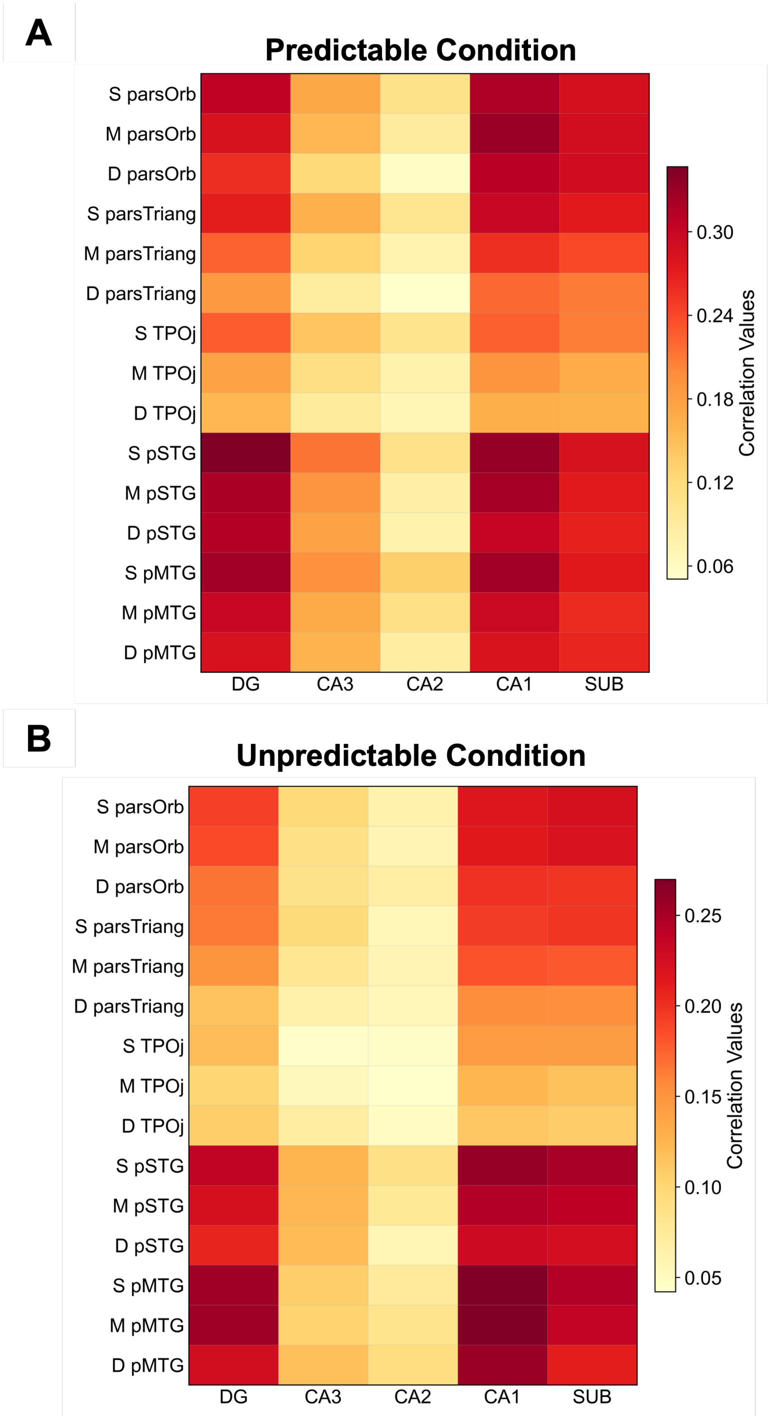


**Supplementary Figure 4: Laminar-level beta series correlation in predictable and unpredictable conditions.** Unthresholded heatmaps of beta series correlation values averaged across subjects for the predictable *(A)* and unpredictable *(B))* conditions. Rows: cortical ROIs across superficial (S), middle (M), and deep (D) layers; columns: hippocampal subfields.

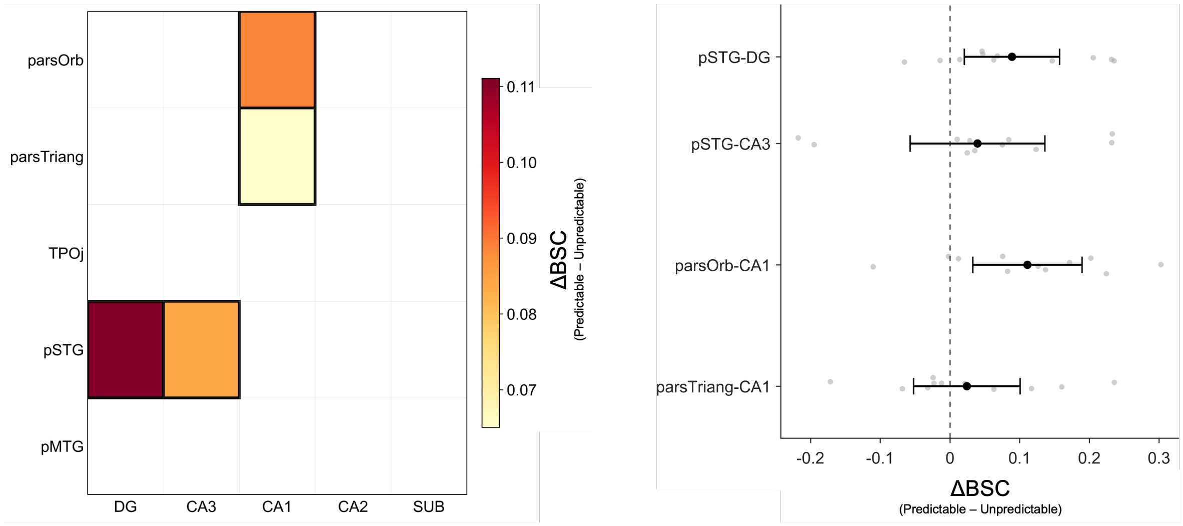


**Supplementary Figure 5: Beta-series connectivity at the level of whole cortical ROIs.** (*A*) Heatmap of group-level significant predictability effects on regional (whole-ROI) BSC connectivity after collapsing across cortical depth. Cells show the mean connectivity difference (predictable - unpredictable) for ROI pairs surviving FDR correction ($q_{\mathrm{FDR}} < 0.05$). Cortical ROIs (pSTG, pMTG, TPOj, pars orbitalis, pars triangularis) and hippocampal subfields (CA1, CA3, DG, SUB) are shown on the two axes. (*B*) Uncertainty of the predictability-related effects. Individual participant values are shown for the difference in beta-series correlation between predictable and unpredictable conditions ($\Delta\mathrm{BSC} = \mathrm{BSC}_{\mathrm{predictable}} - \mathrm{BSC}_{\mathrm{unpredictable}}$). Black dots indicate group means and horizontal error bars the corresponding 95% confidence intervals. Only edges surviving non-parametric sign-flipping permutation testing with FDR correction ($q_{\mathrm{FDR}} < 0.05$) are shown.

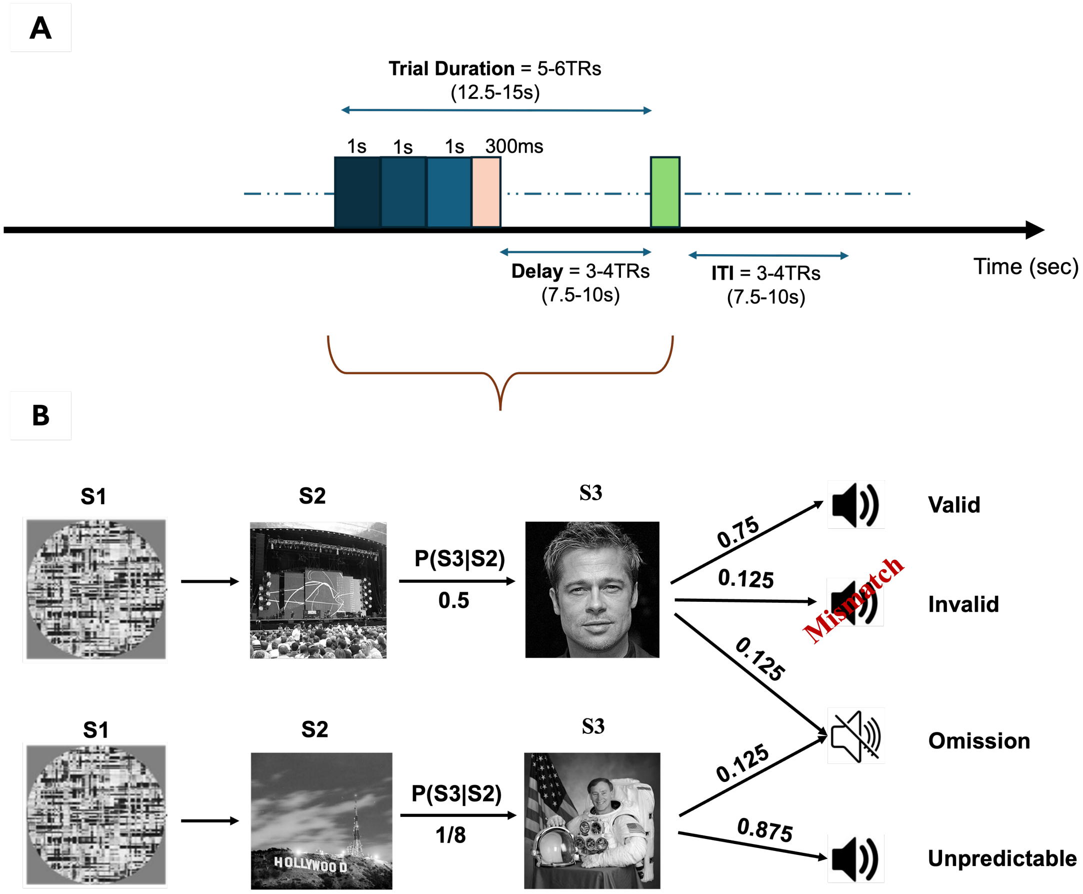


**Supplementary Figure 6: Experimental design.** *(A)* During fMRI acquisition participants listened, in each trial, to an auditory syllable (pink bar, 300 ms) following visual pictures (three pictures presented for one second each). Every trial ended with a cued participant response (green bar) and trials had a random intertrial interval of 3-4 TRs. *(B)* In predictable trials, the auditory syllable could be predicted from the preceding on the basis of a preceding behavioral training. We presented the expected (valid) syllable in 75% of trials, the remaining trials were equally divided between invalid and omission. In unpredictable trials, visual stimuli did not predict the syllable, and omission trials were also included (12.5%).
